# An INF2-dependent actin-mediated step in Inositol 1,4,5-trisphosphate receptor cluster formation and activity

**DOI:** 10.64898/2026.04.09.717539

**Authors:** Maite R. Zavala, Amrapali Ghosh, Suresh K. Joseph, Rajarshi Chakrabarti

## Abstract

Intracellular calcium signaling plays a vital role in regulating various cellular processes including gene regulation, motility, metabolism and cell death. Inositol 1,4,5-trisphosphate receptors (IP3R) on the Endoplasmic Reticulum (ER) are a major cation channel that regulates stimulus-induced calcium release from the ER. While several molecular players regulate activity of IP3R, its regulation by actin filaments were uncharacterized. Here we show that actin filaments polymerized by a specific actin nucleator INF2 facilitates agonist-induced IP3R activity. Our results demonstrate that INF2-mediated actin filaments regulate formation and/or stability of IP3R clusters on the ER that have been previously shown to be hotspots of ER calcium release. Using cell-biological and biochemical techniques we further show that INF2 physically interacts with IP3R isoforms, often at IP3R clusters. While INF2-IP3R interaction is independent of INF2-activity, the ability of INF2 to mediate IP3R clusters is dependent on its actin polymerization activity. Finally, we demonstrate that in addition to its calcium mobilization activity, INF2 on ER specifically regulates IP3R cluster positioning to mediate ER-mitochondrial contacts and facilitate ER to mitochondrial calcium transfer. Overall, these results reveal an actin-dependent step in regulation of IP3R activity both in terms of ER calcium release and modulation of ER-mitochondrial contacts.

**Highlights:**

- INF2-mediated actin filaments potentiate agonist-induced IP3R-mediated ER calcium release without affecting the ER calcium stores per se.
- ER-localization of INF2 is dispensable for its role on IP3R activity. Moreover INF2-mediated actin filaments affect the activity of all IP3R isoforms.
- INF2 interacts with IP3R in an activity and actin filament independent manner through its C-terminal region.
- INF2 regulates IP3R cluster formation in actin-filament dependent manner and thereby regulates IP3R activity.
- Further we show that ER-localized INF2 specifically regulate IP3R cluster positioning thereby promoting ER to mitochondrial contact and calcium transfer.

## Introduction

Calcium (Ca²⁺) ions operating as secondary messengers, play a fundamental role in intracellular signaling regulating various processes like cellular and mitochondrial metabolism ^1,2^, cell motility ^3^, apoptosis ^4^ and vesicular trafficking ^5^ through integration of a complex Ca²⁺ signaling toolkit. Though an increase in intracellular Ca²⁺ levels above the resting 50-100 nM either through external cues ^6^ or spontaneous internal cues ^7,8^ often serve as catalyst to promote Ca²⁺ signaling events, absence of Ca²⁺ spike can also serve as platforms of regulation as shown in certain specific transcriptional repression events ^9^, quiescence ^10^ and autophagy ^11^. Two principal Ca²⁺ sources that cells rely on for signaling events are either Ca^2+^ influx from the extracellular milieu or intracellular Ca^2+^ released by inositol 1,4,5-trisphosphate receptors (IP3Rs) ^12^, ryanodine receptors (RyRs) ^13^, two-pore channels (TPCs) ^14–16^ or the mucolipin subfamily of transient receptor potential channels (TRPML) ^17^. IP3Rs in the non-excitable cells and neurons act as the major Ca²⁺ channel, efflux from which regulate intracellular Ca²⁺ signaling ^18^.

IP3Rs are large-conductance cation channels expressed in most cells that mediate Ca^2+^ release from intracellular stores primarily ER ^19^ and the Golgi apparatus ^20^. IP3Rs are activated by IP3 (generated through phospholipase mediated hydrolysis of plasma membrane lipid phosphatidylinositol 4,5-bisphosphate (PIP2)) and Ca²⁺ ion itself ^21,22^. Three separate genes encode three different isoforms of IP3Rs (IP3R1, IP3R2 and IP3R3) that not only has heterogenous tissue specificity, but also possess distinct activation and deactivation patterns, sensitivity to IP3 and ability to mediate ER to mitochondrial contact (ERMC) and Ca²⁺ transfer ^23^. While IP3Rs have been shown to be arranged in mobile and immobile clusters with Ca^2+^ signaling ^12^ and ERMC ^23^ occurring at these clusters, little is known about the underlying mechanisms that lead to formation of these IP3R clusters. Out of a multitude of factors that have been characterized to regulate IP3R channel activity and function^12^, actin filaments have also been implicated in IP3R-mediated Ca^2+^ release ^24^ from the ER, the mechanism of which still remains uncharacterized.

Specific actin filaments are generated on the ER through the activity of a formin protein INF2 that have been implicated in mitochondrial division ^25^ and ER to mitochondrial Ca²⁺ transfer ^26^. INF2 in cells comes in two isomeric forms: INF2-caax (ER-bound through prenylation of its C-terminal cysteine C1250) and freely cytosol localized INF2-non-caax ^26^. INF2 is activated by raising intracellular Ca^2+^ levels in a Ca^2+^-calmodulin dependent manner ^27–31^ probably relieving the auto-inhibition between its N-terminal region (the DID, Diaphanous Inhibitory Domain) with a C-terminal sequence (the DAD, Diaphanous Autoregulatory Domain) ^27^ which allows it to work both as an actin nucleator and filament elongator. Upon activation, INF2-caax mediated actin filaments specifically on the ER mediates close contacts between ER and mitochondria, facilitating Ca²⁺ transfer from ER to mitochondria that promotes division of the inner mitochondrial membrane (IMM) which is kinetically distinct from Drp1-mediated division of outer mitochondrial membrane (OMM) ^26^. While INF2-mediated actin filaments have been explored more for their role in mitochondrial division process, little is known about its role on ER function.

In this study we demonstrate that INF2-mediated actin filaments regulate agonist-induced IP3R-mediated ER Ca²⁺ release. Acute and chronic depletion of INF2 significantly reduced histamine and carbachol induced ER Ca^2+^ release measured at the level of both cytosolic Ca^2+^ ([Ca^2+^]_C_) and ER Ca^2+^ [Ca^2+^]_ER_ transients. Both isoforms of INF2 could rescue the stimulated Ca^2+^ release in the INF2-depleted cells suggesting that ER localization was dispensable for this function. We further showed that INF2 operates in an IP3R isoform unbiased manner and affects the activity of all the three IP3R isoforms. Through multiple approaches we showed that INF2-caax interacted with IP3R in an activity-independent manner. We further demonstrate that depletion of INF2 significantly reduced assembly of IP3R clusters which could be rescued only by active INF2. Finally, we showed that ER localization of INF2 was imperative to position IP3R2 clusters close to mitochondria which promoted IP3R2-mediated ERMC and Ca²⁺ transfer to mitochondria. These finding suggests that INF2-mediated actin filaments might work to provide a scaffold for regulation of IP3R clusters that not only regulate agonist-induced IP3R activity but also IP3R-mediated ER-mitochondrial crosstalk.

## Results

### INF2 depletion decreases IP3R-mediated ER calcium release

To study the role of INF2 polymerized actin filaments on IP3R-mediated ER Ca^2+^ release, we created CRISPR-mediated INF2-KO in HeLa cells. As shown before, INF2-KO cells (**Figure S1A**) lacked Ca²⁺-mediated “actin burst” (**Figure S1B**). Since the resting cytosolic calcium ([Ca^2+^]_C_) levels showed no difference between the two groups (**Figures S1C and S1D**), we measured agonist-induced cytosolic Ca²⁺ transients using the Cyto-R-Geco calcium probe having a Kd of 0.5 µM. 100 µM histamine induced a significantly higher [Ca^2+^]_C_ transient in the control cells (11.3 ± 3.53-fold) compared to that in the INF2-KO cells (7.8 ± 2.51-fold) (**Figures 1A and 1B**). To test this further we used two sub-maximal doses of histamine (1 µM and 0.1 µM). Similar to the results with the maximal stimulation, both the sub-maximal doses showed significantly higher [Ca^2+^]_C_ transients in the control cells compared to the INF2-KO cells (**Figures 1A and 1B**). Interestingly, 0.1 µM histamine in the control cells yielded a similar [Ca^2+^]_C_ transient (3.8 ± 2.1-fold) as 1µM histamine stimulation in the INF2-KO cells (3.4 ± 1.8-fold). Next, we wanted to test whether this effect was relevant upon acute depletion (siRNA-mediated KD) of INF2 (**Figure S1A**). INF2 KD cells had unchanged resting [Ca^2+^]_C_ (**Figures S1C and S1D**) and showed notably lower [Ca^2+^]_C_ transients compared to the control cells when stimulated by both maximal (100 µM) and sub-maximal (1 µM) doses of histamine (**Figures 1C and 1D**). Interestingly, a significantly higher number of INF2-depleted cells did not respond to the sub-maximal doses of histamine compared to the control cells suggesting that INF2 depleted cells are insensitive to these sub-maximal histamine stimulations (**Figure S1E**). Recording [Ca^2+^]_C_ transients at 1frame/sec revealed that in addition to a decrease in amplitude, INF2 depleted cells not only took longer time to respond but also displayed a slower [Ca^2+^]_C_ transient when stimulated with 1 µM histamine (**Figure 1E**–**1H**). To rule out the possibility that INF2 has a regulatory role on histamine receptors, we stimulated control and INF2-KO cells with 100 µM ATP that uses endogenous purinergic receptors. Similar to histamine stimulations, ATP stimulations also elicited significantly lower [Ca^2+^]_C_ transients in INF2-KO cells (**Figures S1F and S1G**). These results collectively suggest that both acute and chronic depletion of INF2 significantly dampens agonist-induced IP3R-mediated Ca²⁺ release from the ER.

**Figure 1.**
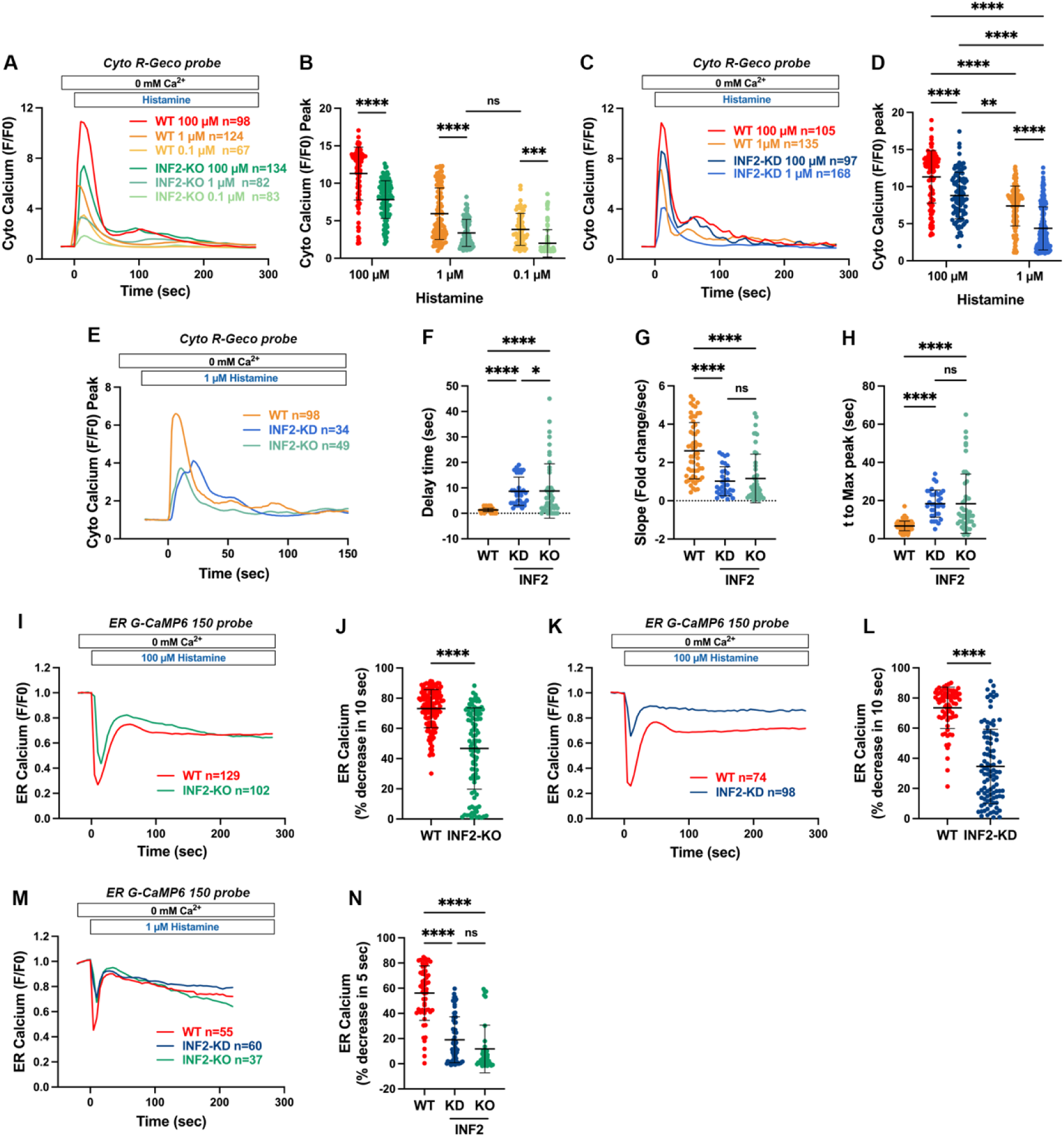
Depletion of INF2 reduces IP3R-mediated calcium release. A: Traces of Cytosolic calcium transients in HeLa WT and INF2-KO cells transfected with Cyto R-GECO plasmid (Kd=0.5 µM), stimulated with different concentrations of histamine (100 µM, 1 µM and 0.1 µM). B: Quantification of maximal peak of cytosolic Ca2+ release as shown in A, expressed as fold change F/F0. Data from 80-130 cells across 3 - 4 independent experiments. Graphs shows Mean ± S.D. Two-way ANOVA used. **** p <0.0001, *** p=0.0005, * p=0.0133. C: Traces of Cytosolic calcium transients in HeLa WT and INF2-KD cells transfected with Cyto R-GECO plasmid (Kd=0.5 µM), stimulated with different concentrations of histamine (100 µM and 1 µM). D: Quantification of maximal peak of cytosolic Ca2+ release as shown in C, expressed as fold change F/F0. Data from 90-160 cells across 3 - 4 independent experiments. Graphs shows Mean ± S.D. **** p <0.0001, ** p=0.0034. E: Traces of Cytosolic calcium transients in HeLa WT, INF2-KD and INF2-KO HeLa cells transfected with Cyto R-GECO plasmid (Kd=0.5 µM), stimulated with 1 µM histamine and recorded at 1 frame/sec. F: Quantification of the time delay between histamine stimulation onset of cytosolic calcium transients in WT, INF2 KD and INF2 KO HeLa cells imaged in Fig 1E. Data from 58 (WT), 32 (KD) and 48 (KD) cells across 3 independent experiments in dataset shown in 1E. Graph shown Mean ± SD. **** p <0.0001, * p=0.0295. G: Quantification of slope of cytoplasmic calcium rise, expressed as fold change/sec from Dataset shown in Fig 1E in WT, INF2 KD and INF2 KO HeLa cells. Data from 58 (WT), 32 (KD) and 48 (KD) cells across 3 independent experiments. Graph shown Mean ± SD. **** p <0.0001 H: Quantification of time taken for cytoplasmic calcium spike to reach the max value from Dataset shown in Fig 1E in WT, INF2 KD and INF2 KO HeLa cells. Data from 58 (WT), 32 (KD) and 48 (KD) cells across 3 independent experiments. Graph shown Mean ± SD. **** p <0.0001 E: Traces of ER calcium transients in HeLa WT and INF2-KO cells transfected with ER-GCaMP6-150 plasmid (Kd=150 µM), stimulated with histamine (100 µM). F: Quantification of ER calcium expressed as % of decrease in 10 sec from traces shown in E. Data from 129 (WT) and 102 (INF2-KO) cells from 3 independent experiments in each case. Graphs shows Mean ± S.D. unpaired Student’s t-test used. **** p<0.0001. G: Traces of ER calcium transients in HeLa WT and INF2-KD cells transfected with ER-GCaMP6-150 plasmid (Kd=150 µM), stimulated with histamine (100 µM). H: Quantification of ER calcium expressed as % of decrease in 10 sec from traces shown in G. Data from 74 cells (WT) and 98 cells (INF2-KD) across 3 independent experiments. Unpaired student’s t-test used. **** p<0.0001. I: Traces of ER calcium transients in HeLa WT, INF2-KO and INF2-KD cells transfected with ER G-CaMP6-150 plasmid (Kd=150 µM), stimulated with1uM histamine treatment. J: Quantification of ER calcium release from I, expressed as % of decrease in 5 sec. Data from 55 (WT), 60 (INF2-KD) and 37 (INF2-KO) cells from 3 independent experiments. Graphs shows Mean ± S.D. Significant was measured using One-way ANOVA. **** p<0.0001.

So far, we evaluated ER Ca²⁺ release through measuring [Ca^2+^]_C_ transients, and we wanted to directly evaluate kinetics of ER calcium ([Ca^2+^]_ER_) release in control and INF2 depleted cells. Control and INF2 KO (**Figures 1I-1J**; **1M-1N**) or INF2 KD (**Figures 1K-1L**) cells were transfected with the high affinity luminal ER-GCamp150 probe and stimulated with either 100 µM histamine (**Figures 1I-1L**) or 1 µM histamine (**Figures 1M-1N**). While control cells show about 73% acute decrease in ER Ca^2+^ content upon 100 µM histamine stimulation (**Figures 1I-1J**), both INF2-KO cells (46.7 ± 26 %) and INF2-KD cells (34.6 ± 24 %) showed significantly lower Ca^2+^ release from ER (**Figures 1I-1L**). A similar trend was noted for the sub-maximal stimulation where control cells showed 56.1 ± 21% decrease in ER Ca^2+^ upon stimulation, significantly higher than both INF2-KO (11.7 ± 18.9%) and INF2-KD (18.9 ± 18.2 %) cells (**Figures 1M-1N**). We explored this further using a low-sensitive ER-LAR-Geco Ca^2+^ probe having a Kd of 24 µM. Using this probe the control cells showed 28.9 ± 9.5 % acute [Ca^2+^]_ER_ release which was significantly stronger than both INF2-KO (1.78 ± 3.5%) and INF2-KD (5.47 ± 4.7%) cells (**Figures S1H and S1I**). We also measured the abundance of ER Ca²⁺ stores through 2µM Thapsigargin treatments which showed similar [Ca^2+^]_C_ transients among the control and INF2-depleted (KO and KD) cells signifying that the ER stores were not pre-exhausted upon INF2 depletion (**Figure S1C, S1J**). Cumulatively, these results confirm that both acute and chronic depletion of INF2 significantly subdue agonist-induced IP3R-mediated ER Ca²⁺ release measured at the level of cytosolic and ER Ca²⁺ transients.

### ER localization of INF2 is dispensable for IP3R-mediated calcium release

In cells, INF2 is expressed in two isomeric forms: ER-bound INF2-caax and cytosolic INF2-non-caax. We next assessed whether ER residency of INF2 was necessary for regulation ER Ca^2+^ release. We re-expressed GFP-INF2-caax (**Figures 2A and 2B; S2A**) and GFP-INF2-noncaax (**Figures 2C and 2D; Figure S2A**) in HeLa INF2-KO cells and measured [Ca^2+^]_C_ transients upon 100 µM histamine stimulation. While INF2-caax (ER) completely rescued histamine-induced [Ca^2+^]_C_ transients in the INF2-KO cells (**Figures 2A and 2B**), INF2-noncaax (cyto) also could significantly salvage [Ca^2+^]_C_ transients in INF2-KO cells (**Figures 2C and 2D**). Further, we measured [Ca^2+^]_ER_ transients using the low affinity ER LAR-Geco probe following re-expression of INF2-caax or INF2-noncaax. Re-expression of both INF2 isoforms in INF2-KO cells also rescued the [Ca^2+^]_ER_ transients further confirming that ER residency of INF2 was not required to mediate IP3R-mediated ER Ca²⁺ release (**Figures 2E and 2F**). Interestingly we noticed that while baseline resting [Ca^2+^]_C_ in INF2-rescue cells remained unaltered compared to the control cells (**Figures S1C and S1D**), 2 µM Thapsigargin induced [Ca^2+^]_C_ transients revealed slightly depleted ER stores (**Figures S1C and S1J**). However, the mildly depleted ER stores in the INF2-rescue conditions did not alter histamine-induced [Ca^2+^]_C_ or [Ca^2+^]_ER_ transients. Though INF2 in cells is mostly auto-inhibited due to the intrinsic DID (N-term)-DAD (C-term) binding and therefore tightly regulated, we still wanted to evaluate whether overexpression of INF2-isoforms positively regulated histamine-induced IP3R-mediated [Ca^2+^]_ER_ transients. To do this, we transfected either INF2-caax or INF2-non-caax in HeLa WT cells (**Figure S2B**) and measured histamine-induced [Ca^2+^]_C_ or [Ca^2+^]_ER_ transients using low-affinity probes Cyto-LAR-Geco (Kd 12 uM) (**Figures 2G and 2H**) and ER-LAR-Geco (Kd 24 uM) (**Figures 2I and 2J**) respectively. Overexpression of INF2 isoforms significantly enhanced both [Ca^2+^]_C_ or [Ca^2+^]_ER_ transients compared to the un-transfected control cells but did not alter the resting [Ca^2+^]_C_ or [Ca^2+^]_ER_ levels (**Figures S2C-S2E**). These results suggest that abundance of either of the INF2 isoforms significantly regulate histamine-induced IP3R-mediated [Ca^2+^]_C_ and [Ca^2+^]_ER_ transients in HeLa cells.

**Figure 2.**
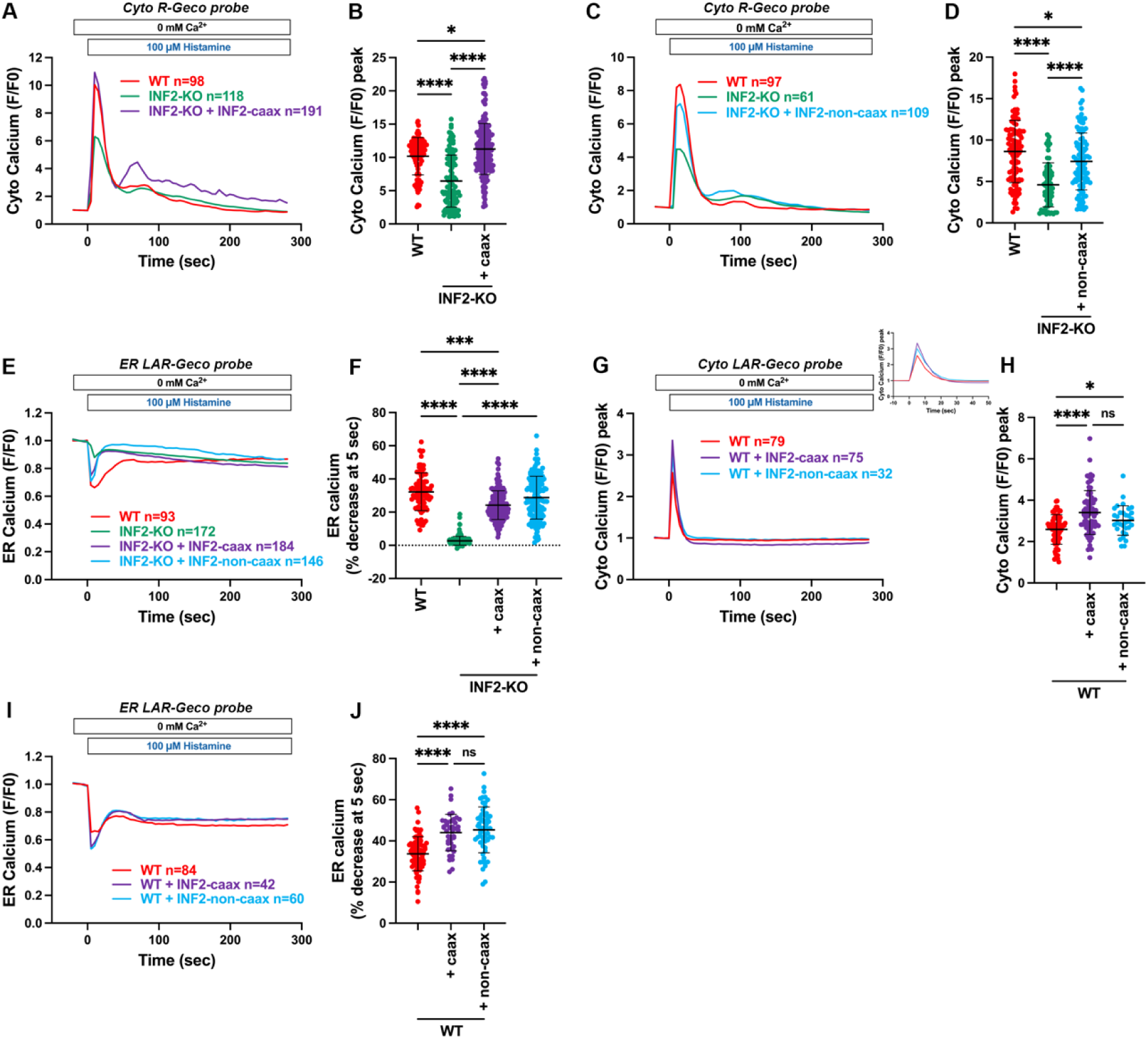
ER localization of INF2 is dispensable for IP3R-mediated calcium release. A: Traces of Cytosolic calcium transients in HeLa WT, INF2-KO and INF2-caax rescued cells transfected with Cyto R-GECO plasmid (Kd=0.5 µM), stimulated with 100 µM histamine. B: Quantification of maximal peak of cytosolic Ca2+ release as shown in A, expressed as fold change F/F0. Data from 98 (WT), 118 (INF2-KO) and 191 (IN2-KO + INF2-caax) cells across 3 - 4 independent experiments. One way ANOVA used. **** p<0.0001 * p=0.0447 C: Traces of Cytosolic calcium transients in HeLa WT, INF2-KO and INF2-noncaax rescued cells transfected with Cyto R-GECO plasmid (Kd=0.5 µM), stimulated with 100 µM histamine. D: Quantification of maximal peak of cytosolic Ca2+ release as shown in C, expressed as fold change F/F0. Data from 97 (WT), 61 (INF2-KO) and 109 (IN2-KO + INF2-noncaax) cells across 3 independent experiments. One way ANOVA used. **** p<0.0001 * p=0.0318 E: Traces of ER calcium transients in HeLa WT, INF2-KO and INF2-KO rescue cells with INF2-Caax or INF2–noncaax co-transfected with ER-LAR-GECO (Kd=24 µM), stimulated with 100 μM Histamine treatment. F: Graph quantifying changes in ER calcium release upon the Histamine treatment, expressed as % of depletion at 5 sec. Data from 93 (WT), 172 (INF2-KO), 184 (INF2-KO + INF2-caax) and 146 (INF2-KO + INF2-noncaax) cells across 3 independent experiments. Graph shows Mean ± S.D. One-way ANOVA used. **** p<0.0001, *** p=0.0004. G: Traces of cytosolic transients in HeLa WT cells either alone or over-expressing INF2-caax or INF2-noncaax co-transfected with Cyto LAR-GECO (Kd=12 µM) probe, stimulated with 100 µM histamine H: Quantification of maximal peak of cytosolic calcium from G, expressed as F/F0. Data from 79 (WT), 75 (WT + INF2-caax) and 32 (WT + INF2-noncaax) cells across 3 independent experiments Graphs shows Mean ± S.D. Significant was measured using One-way ANOVA. **** p<0.0001, *p=0.0483. I: Traces of ER calcium transients in HeLa WT cells either alone or over-expressing INF2-caax or INF2-noncaax co-transfected with ER-LAR-GECO probe (Kd=24 µM), stimulated with after 100 µM histamine. J: Quantification of ER calcium release from I, expressed as % of decrease in 5 sec. Data from 84 (WT), 42 (WT + INF2-caax) and 60 (WT + INF2-noncaax) cells across 3 independent experiments Graph shows Mean ± S.D. Significant was measured using One-way ANOVA. **** p<0.0001

### Depletion of INF2 affects the function of all IP3R isoforms

Since HeLa cells express all the three isoforms of IP3R (IP3R1, IP3R2 and IP3R3), it is unclear whether INF2 has a functional bias towards any of the IP3R isoforms. To address this, we utilized previously characterized HEK-293 cell lines expressing either IP3R1 (IP3R2/3 DKO) or IP3R2 (IP3R1/3 DKO) or IP3R3 (IP3R1/2 DKO) ^32^. Firstly, we performed acute knockdown of INF2 in these cell lines (**Figure S3A**), stimulated them with 100 µM carbachol in Ca^2+^ free extracellular medium and recorded [Ca^2+^]_C_ transients. Acute depletion of INF2 significantly reduced [Ca^2+^]_C_ transients mediated by either IP3R1 (**Figures S3B and S3C**) or IP3R2 (**Figures S3D and S3E**) or IP3R3 (**Figures S3F and S3G**). Further, we created CRISPR-INF2-KO in either IP3R2 (IP3R1/3 DKO) or IP3R3 (IP3R1/2 DKO) expressing HEK cell lines and validated them for the absence of INF2 (**Figure S3A**). Similar to results obtained upon acute depletion, INF2 KO in either IP3R2 (IP3R1/3 DKO) (**Figures S3D and S3E**) or IP3R3 (IP3R1/2 DKO) (**Figures S3F and S3G**) expressing cell lines significantly dampened carbachol induced [Ca^2+^]_C_ transients. Similar to results obtained using HeLa cells, re-introduction of either of the INF2 isoforms completely rescued both [Ca^2+^]_C_ (**Figures S3H and S3I**) and [Ca^2+^]_ER_ (**Fig. S3J and S3K**) transients in INF2 depleted IP3R2 (IP3R1/3 DKO) HEK cells. Collectively these results confirm that loss of INF2 affects [Ca^2+^]_C_ and [Ca^2+^]_ER_ transients induced by all IP3R isoforms individually.

### INF2 interacts with IP3R-isoforms independent of its actin polymerization activity

We wanted to test whether regulation of IP3R function by INF2 is mediated through a direct interaction or is an effect of a signaling cascade. Proximity Ligation Assay (PLA) against IP3R2/INF2 and IP3R3/INF2 using validated IP3R2 and IP3R3 antibodies (**Figure S4A**) in HeLa control and INF2-KO cells revealed significant proximity of both IP3R2 (**Figures 3A and 3B**) and IP3R3 (**Figures S4B and S4C**) with INF2 in control cells which was significantly reduced in INF2-KO cells. Re-introduction of INF2 isoforms in INF2-KO cells significantly rescued the density of PLA dots for both IP3R2 (**Figures 3A and 3B**) and IP3R3 (**Figures S4B and S4C**). Interestingly there was significantly increased density of PLA dots between IP3R3- INF2-caax compared to IP3R3-INF2-non-caax suggesting that INF2-caax on the ER might preferentially interact better with IP3R3 (**Figures S4B and S4C**). This phenotype was not relevant for IP3R2-INF2 interactions (**Figure 3A and 3B**).

**Figure 3:**
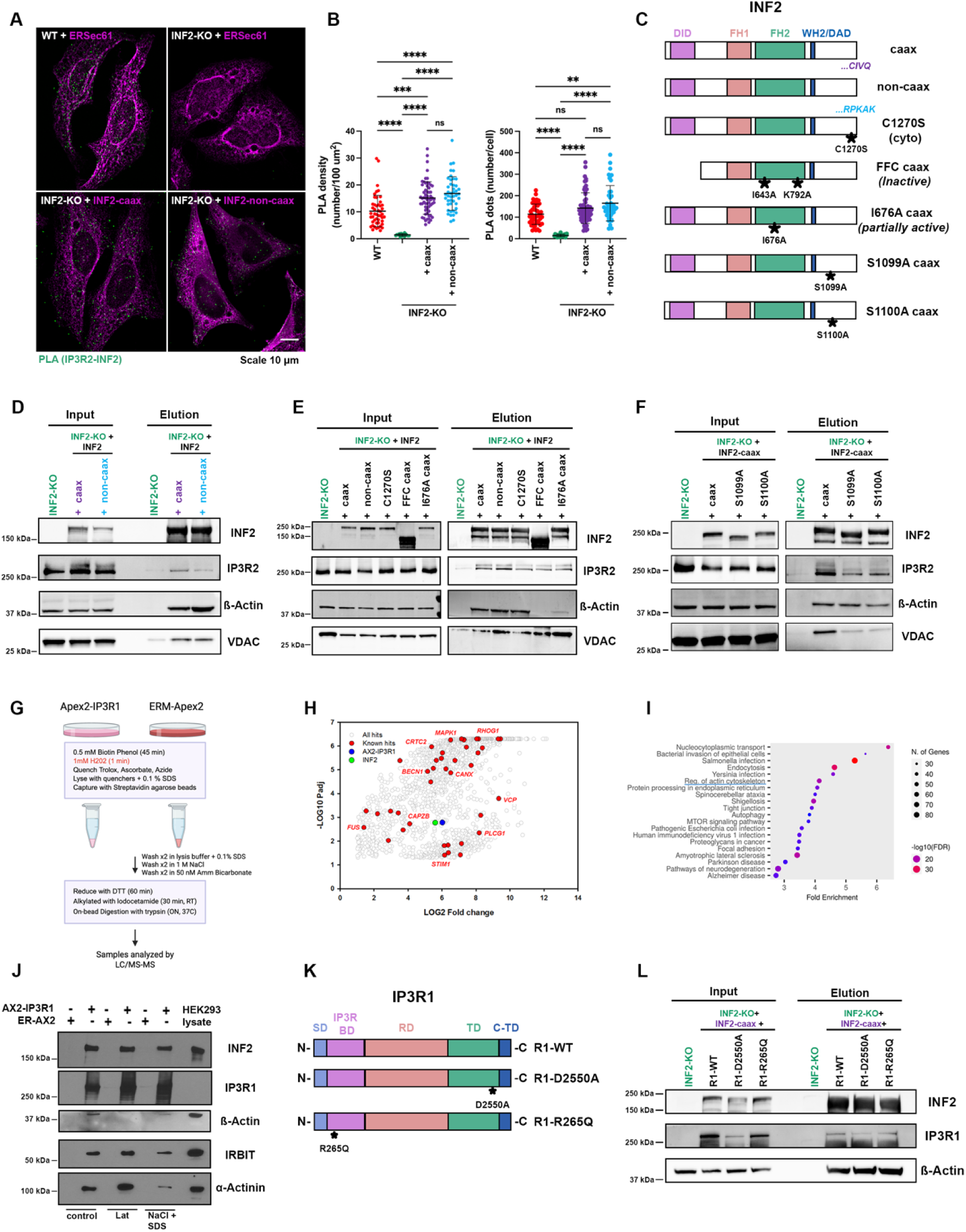
INF2 interacts with IP3R-isoforms independent of its actin polymerization activity. A: Representative fixed-cell images from Proximity Ligation Assay (PLA) against IP3R2 and INF2 in HeLa WT, INF2-KO and INF2-KO rescue cells. WT and INF2-KO cells were transfected with GFP-ERSec61β to label ER (magenta) prior to PLA staining. PLA foci shown in green. Scale: 10 µm. B: Quantification of PLA density expressed as number of dots per 100 µm2 (left), and PLA dots, expressed as number/cell (right). Data from 48 (WT), 20 (INF2-KO), 60 (INF2-KO+INF2-caax) and 44 (INF2-KO+INF2-noncaax) cells from 3 independent experiments Graph shows Mean ± S.D. One-way ANOVA used. **** p<0.0001, *** p=0.0001 (PLA density, left); **** p<0.0001, ** p=0.0012 (PLA dots, right). C: Illustration of INF2 domains structure, showing the WT and with different mutations (C1270S-caax: cytosolic localization; FFC caax (inactive): ΔDID/ I643A/K792A and I676A caax (partially active). References: DID, Diaphanous Inhibitory Domain; FH1, Formin Homology 1; FH2, Formin Homology 1, WH2/DAD, Diaphanous Autoregulatory Domain. D: Western blot images of candidate proteins from GFP-trap pull-down assay, in HEK IP3R2 expressing INF2-KO cells transfected with GFP-tagged INF2-Caax or INF2-noncaax. E: Western blot images of candidate proteins from GFP-trap pull-down assay, in HEK IP3R2 expressing INF2-KO cells transfected with GFP-tagged INF2 constructs (WT and mutants). F: Western blot images of candidate proteins from GFP-trap pull-down assay, in HEK IP3R2 expressing INF2-KO cells transfected with GFP-tagged INF2 constructs (WT and serine-mutants). G: Workflow of APEX2-labelling biotinylating assay using mass spectroscopy, to identify IP3R1 proximity partners in HEK-IP3R TKO cells transfected with either APEX2-IP3R1 or APEX2-ER (used as control) constructs. H: Plot of known interactors of IPR31, showing INF2. Fold change was calculated based on enrichment over control (ERM-APEX2). An ‘imputed’ averaged value from all samples was used to calculate fold enrichment. This accounts for the plateau in the Y-axis. I: Graph showing significant enrichment of proteins involved in regulation of actin cytoskeleton. J: Representative western blot images from streptavidin pull-downs following APEX2 labelling as shown in assay in panel G in different conditions. K: Illustration of IP3R domains showing the two different mutations: IP3R1-D2550A pore dead mutant and IP3R1-R265Q, ligand binding motif mutant. References: SD, Suppressor domain; IP3R BD, IP3R binding domain; RD, Regulatory domain; TD, Transmembrane domain; C-TD: Carboxy-terminal domain. L: Western blot images of candidate proteins from GFP-trap pull-down assay performed in HEK IP3R-TKO-INF2-KO cells re-expressed with IP3R1 (WT/mutants) and GFP-INF2-caax.

To further validate this interaction, we carried out pull-down assays using GFP-trap agarose beads to confirm whether endogenous IP3R2 physically interacts with INF2. GFP-tagged INF2 isoforms (caax and non-caax) (**Figure 3C**) were transfected into IP3R2 expressing-INF2-KO HEK cells for 24 hours, lysed and pulled down using GFP-trap beads. Endogenous IP3R2 was detected in the eluates of both INF2-caax and INF2-non-caax (albeit at lower abundance) suggesting an interaction between the two proteins. We also detected β-actin (positive control) and IP3R interactor VDAC in the eluates of both INF2 isoforms (**Figure 3D**). Since we detected a significant amount of β-actin in the eluates, we wondered whether IP3R2 interactions with INF2 were mediated actin filaments. To test this, cells expressing GFP-INF2 isoforms (caax and non-caax) were treated with 0.5 µM latrunculin A (Lat A) for 15 mins prior to lysis and GFP-trap pull-down. Interestingly, while IP3R2 interactions with INF2-caax was not affected by actin depolymerization, interaction with INF2-non-caax was significantly reduced (**Figure S4D**). Similar fate was also observed for VDAC (**Figure S4D**). However, we still detected a significant amount of β-actin in the pull-down eluates which might be actin monomers being brought down by INF2 isoforms. To further test the relevance of actin filaments in IP3R2-INF2 interactions, we used two mutant INF2-caax constructs: (1) N-term DID domain truncated FFC construct bearing two mutations, I643A and K792A, in the FH2 domain that makes it inactive; (2) a full-length INF2-caax bearing I676A mutation that makes it partially active (**Figures 3C; S1B**). While the FFC-caax mutant did not interact with β-actin, I676A mutant had significantly lower β-actin in the pull-down eluates (**Figure 3E**). IP3R2 was detected in the eluates of both these constructs at levels similar to those from INF2-caax (WT), INF2-non-caax and a cytosol localized INF2-caax C1270S mutant (**Figure 3E**). These results further strengthened the fact that IP3R2-INF2 interaction was independent of actin filaments. INF2 was recently shown to be phosphorylated at the C-terminal (S1077 (human)/ S1099 (mouse)) by AMPK that regulates mitochondrial fission ^33^. Moreover, it was also shown to interact with another ER integral membrane protein, MAL, through the C-terminal 200 aa ^34^ suggesting that the C-terminal region might be an important interacting hub for INF2. To identify whether the C-terminal Serine was important for its interaction with IP3R, we created an INF2- caax bearing S1099A and S1100A mutations (**Figure 3C**). Expressing them in INF2-KO cells and performing the GFP-trap assay revealed that both the mutants tested was significantly impaired for its interaction with IP3R2 (**Figures 3F**; **S4E**). Thus, it is evident that either the phosphorylation at S1099/1100 or the serine residues *per se* at the C-terminal of INF2 facilitate its binding with IP3R2. Collectively these results confirm an actin-independent interaction between INF2-caax and endogenous IP3R2.

Taking a more unbiased approach, we performed an APEX2 labelling assay to identify binding partners for IP3R1 in HEK-293 cells. APEX2-IP3R1 and an APEX2-ER construct was transfected into HEK-293-TKO (IP3R1/2/3 KO) cells, incubated with biotin, pulsed with 1 mM H_2_O_2_ for 1 min, quenched, lysed and pulled-down with streptavidin agarose beads. The precipitated molecules were analyzed using LC-MS/MS (**Figure 3G**). Several known interactors were identified and satisfyingly endogenous INF2 emerged as a potent interactor of IP3R1 (**Figure 3H; Dataset 1**). Additional cytoskeletal proteins identified include Arp2/3 components mDia 1/2, Spectrin, α-actinin and Filamin A (**Dataset 1**). In fact, further analysis revealed a significant enrichment of proteins involved in regulation of the actin cytoskeleton (**Figure 3I**). We validated some of these targets. INF2, IRBIT and α-actinin were validated in immunoblots both in control and latrunculin A treated cells suggesting an actin-filament independent binding of these molecules with IP3R1 (**Figure 3J**). While a high salt elution weakened the abundance of IRBIT and α-actinin in the eluate, the abundance of INF2 was not affected, suggesting a rather strong interaction with IP3R1 (**Figure 3J**). Finally, we tested the ability of IP3R1 mutants compromised for IP3-binding (R1-R265Q) or channel function (R1-D2550A) to interact with INF2. WT-IP3R1 and the mutants (**Figure 3K**) were transfected into IP3R TKO INF2-KO HEK cells along with GFP-INF2-caax. GFP-trap precipitation showed that the IP3-binding and pore forming ability is dispensable for IP3R1 binding with INF2 (**Figure 3L**). Therefore, we conclude through various approaches that INF2-caax interacts with IP3R isoforms in an actin-filament independent way which also does not require IP3R activity.

### INF2 regulates formation of functionally competent IP3R clusters in cells

Since INF2 depletion significantly affected agonist-induced (histamine, ATP and carbachol) IP3R-mediated Ca^2+^ transients, we wanted to decipher the molecular mechanism underlying this phenotype. Possible mechanism(s) include insufficient production of agonist stimulated IP3 molecules, altered abundance of IP3R on the ER or changes in IP3R organization on the ER. To test out the first possibility, we transfected YFP-tagged PH-PLC-δ construct ^35^ that binds to PIP2 at the plasma membrane. Agonist-induced hydrolysis of PIP2 and IP3 production allows re-localization of the probe to cytosol which we use as a surrogate to measure IP3 production. 100 µM histamine and carbachol treatments of control and INF2-KO HeLa (**Figures S5A and S5B**) and IP3R2-HEK cells (**Figures S5A and S5C**) respectively, showed similar kinetics of the probe re-localization indicating that the abundance and kinetics of IP3 production might not be affected by INF2 depletion. Next, we looked at IP3R abundance upon INF2 depletion. While IP3R2 abundance seemed to increase slightly in HeLa INF2-KO cells, abundance of IP3R1 and IP3R3 remained unchanged in the INF2-depleted cells (**Figure 4A**). Similarly, IP3R2 (**Figure 4B**) and IP3R3 (**Figure S5D**) abundance in IP3R2 expressing-INF2-KO HEK cells and IP3R3 expressing-INF2-KO HEK cells respectively were also unchanged compared to their respective control cells. These results further shows that chronic depletion of INF2 does not affect the abundance of IP3R isoforms in either HeLa or HEK cells.

**Figure 4.**
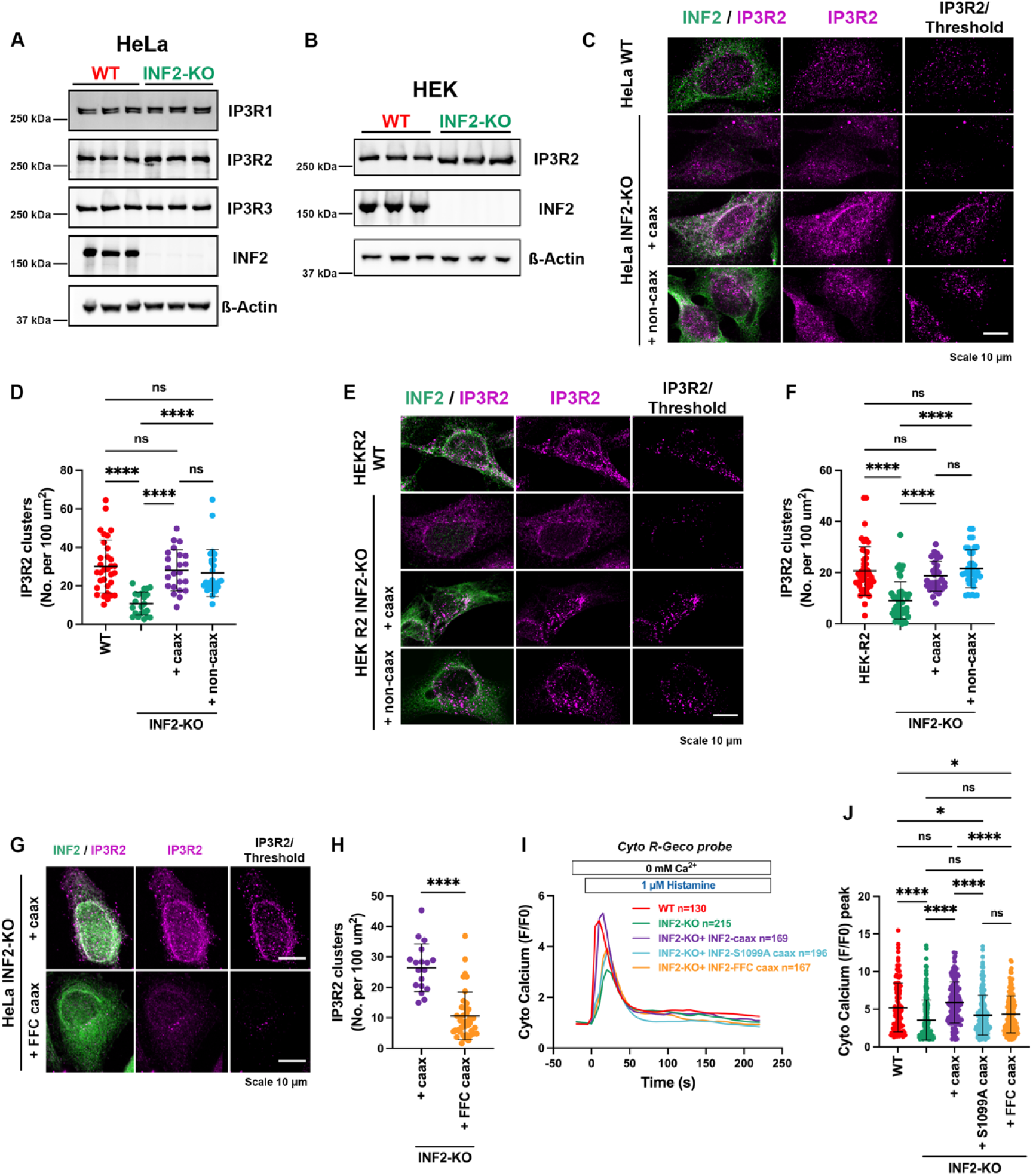
INF2 regulates formation of functionally competent IP3R clusters in cells. A: Western blots showing the expression of all IP3R isoforms in WT and INF2-KO HeLa cells. β-Actin was used as loading control. B: Western blots showing expression of IP3R2 in HEK IP3R2 expressing (R1/R3 DKO) WT or INF2-KO. C: Representative confocal micrographs of HeLa WT, INF2-KO and INF2-KO rescue cells (transfected with GFP-INF2-Caax or GFP-INF2-non-caax), immunoassayed for INF2 (green) and IP3R2 (magenta). IP3R2 clusters after thresholding shown in the right-most panel. Scale: 10 µm. D: Quantification of IP3R2 clusters, expressed as number of clusters per 100 µm2 of cell area (from dataset in C). Data from 32 (WT), 22 (INF2-KO), 23 (INF2-KO+INF2-caax) and 27 (INF2-KO+INF2-noncaax) from 2 independent experiments. E: Representative confocal micrographs of HEK-IP3R2 expressing cells (WT, INF2-KO and rescues), immunoassayed for INF2 (green) and IP3R2 (magenta). IP3R2 clusters after thresholding shown in the right-most panel. Scale: 10 µm. F: Quantification of IP3R2 clusters, expressed as number of clusters per 100 µm2 of cell area (from data set in E). Data from 47 (WT), 49 (INF2-KO), 32 (INF2-KO+INF2-caax) and 40 (INF2-KO+INF2-non-caax) cells from 2 independent experiments Graphs shows Mean ± S.D. Significant was measured using One-way ANOVA. **** p<0.0001. G: Representative confocal micrographs fixed-HEK IP3R2 expressing INF2-KO cells transfected with either GFP-INF2-Caax (WT) or GFP-INF2 FFC caax (inactive), immunoassayed for IP3R2 (magenta). Scale: 10 µm. H: Quantification of IP3R2 clusters, expressed as number of clusters per 100 µm2 (from data set in G). Data of 18 cells for (INF2-caax-WT) and 39 cells (FFC caax-inactive) from 2 independent experiments. Graphs show Mean ± S.D. unpaired Student’s t-test used. **** p<0.0001. I: Cytosolic Ca2+ transients induced by 1 µM histamine in Cyto R-GECO transfected HeLa WT (red), INF2-KO (green), INF2-KO + INF2-caax WT (purple), INF2-FFC-caax inactive mutant (orange) or INF2-caax-S1099A binding mutant (yellow). J: Quantification of maximal peak of Cyto Ca2+ from I, expressed as F/F0. Data from 130 (WT), 215 (INF2-KO), 169 (INF2-KO + INF2 caax), 167 (INF2-KO + INF2-FFC-caax) and 196 (INF2-KO + INF2-caax S1099A) cells across 5 independent experiments. Graphs show Mean ± S.D. One-way Anova used. **** p<0.0001, * p= 0.0427 (WT vs. + FFC caax), * p=0.0110 (WT vs. + S1099A).

Next, we performed immunofluorescence staining of IP3R2 in cells. Fixed cell analysis of IP3R2 and IP3R3 in HEK cells showed a faint homogenous staining of endogenous IP3R isoforms (**Figure S4A**) on the ER with intermittent bright immunoreactive cluster also localized to the ER (**Figure S4A**). Immunoreactive signals for both IP3R2 and IP3R3 was not detected outside the ER signal (**Figure S4A**). Surprisingly depletion of INF2 drastically decreased the density of the bright immunoreactive cluster of endogenous IP3R2 in HeLa cells (**Figure 4C and 4D**) and in IP3R2-expressing HEK cells (**Figure 4E and 4F**). We further stained for endogenous IP3R3 in HeLa cells and saw a similar reduction in IP3R3 clusters upon depletion of INF2 (**Figures S5E and S5F**). Further we stained for endogenous IP3R3 and simultaneously performed PLA for IP3R3/ INF2 which revealed that about 63.2 ± 7.9% of the PLA dots are on IP3R3 clusters suggesting that substantial interaction between IP3R3 and INF2 occurred at IP3R3 clusters (**Figures S5G and S5H**). Since both isoforms of INF2 could significantly rescue agonist induced [Ca^2+^]_C_ and [Ca^2+^]_ER_ transients in HeLa and HEK cells, we wanted to see whether re-introduction of INF2 isoforms also rescued IP3R2 clustering. Satisfyingly both INF2 isoforms could completely recover clustering of endogenous IP3R2 in both the cell lines (**Figures 4C-4F**). While INF2-activity seems dispensable for its ability to bind IP3R, we wanted to test whether activity is required for IP3R clustering and agonist-induced IP3R activity. While re-introduction of INF2-caax (WT) rescued both IP3R2 cluster formation (**Figures 4G and 4H**) and histamine-induced [Ca^2+^]_C_ transients in HeLa INF2-KO cells (**Figures 4I and 4J**), the inactive FFC-caax mutant, which still interacts with IP3R2 (**Figure 3E**) failed to significantly rescue either of the phenotypes (**Figures 4G-4J**). Additionally, the INF2-caax S1099A mutant, with reduced IP3R2 binding, was also severely compromised for its ability to rescue histamine-induced [Ca^2+^]_C_ transients in INF2-KO cells (**Figures 4I-4J**). Interestingly INF2-noncaax S1099E mutant could significantly restore [Ca^2+^]_C_ transients in INF2-KO cells similar to WT-INF2-noncaax indicating the importance of phosphorylation at this specific site for agonist-induced ER Ca^2+^ release (**Figure S5I and S5J**). These results suggest that while INF2-IP3R2 interactions are independent of INF2-activity, IP3R2 cluster formation and agonist-mediated IP3R activity is dependent on INF2-activity. We rule out INF2-mediated ER structural aberration as a plausible mechanism as this ER phenotype is exclusively rescued by INF2-caax (**Figure S5K**) while IP3R activity and cluster assembly is rescued by both INF2 isoforms. Collectively these results suggest that INF2 plays a significant role in formation and/or stability of endogenous IP3R clusters without altering its absolute abundance on ER. Our results indicated that both IP3R2 and IP3R3 loses their clustering ability upon depletion of INF2, a phenotype that is completely rescued by re-introduction of INF2. Previous results have shown that Ca^2+^ release preferentially occur close to the immobile IP3R clusters on the ER ^36^ and we propose that cluster formation might be regulated by INF2.

### INF2-caax specifically regulate ER-Mito contacts and ER to mito calcium transfer

Our results show that both INF2-caax on ER and INF2-non-caax in the cytosol can interact with IP3Rs and regulate cluster formation thereby allowing for agonist-induced ER Ca²⁺ release. However, we had previously shown that ER localization of INF2 is indispensable for its role in ER-Mitochondrial Ca²⁺ transport and mitochondrial division ^26^. Therefore, we reasoned that INF2-caax has unique roles beyond IP3R clustering to mediate ER-mitochondrial tethering and agonist-induced ER to mitochondrial Ca²⁺ transfer. To this end we imaged for GFP-Tom20 and endogenous IP3R2 in INF2-KO cells rescued with either INF2-caax or INF2-non-caax. While in both cases we noticed IP3R2 cluster formation, finer evaluation of the clusters revealed that in the case of INF2-caax re-expression the IP3R2 clusters were more aligned with mitochondria compared to the ones with INF2-non-caax re-expression (**Figures S6A and S6B**). We therefore wanted to evaluate whether depletion of INF2 had any profound effect on the ability of IP3R2 to mediate ER-Mito contacts (ERMC). We performed PLA using VAP-B (ER) and Tom20 (Mito) as a surrogate for ERMC. While IP3R2 expressing HEK cells showed an increased density of PLA dots compared to IP3R-TKO HEK cells (**Figures 5A and 5B**), depletion of INF2 from these IP3R2-expressing cells significantly diminished the ability of IP3R2 to create ER-Mito contacts (**Figures 5A and 5B**). Finer evaluation of these contacts from thin-section TEM images showed that depletion of INF2 in the IP3R2 expressing cells compromised the tight contacts (0-20 nm) between ER and mitochondria (**Figures 5C and 5D**). Infact chronic depletion of INF2 in HeLa cells (that has all the IP3R isoforms) also drastically reduces close contacts and increases the minimum distance between ER and mitochondria as evaluated from thin-section TEM analysis (**Figures S6C-S6E**). Re-introduction of INF2- caax completely rescued the density of PLA dots (VAP-B/Tom20) but INF2-non-caax could only partially rescue the phenotype (**Figures 5A and 5B**). The abundance of VAP-B, Tom20 and PTPIP51 was similar across all the conditions tested (**Figure S6F**). Further we functionally validated the relevance of INF2-mediated ERMC through recording mitochondrial Ca²⁺ uptake in IP3R2-expressing HEK cells. Depletion of INF2 in IP3R2-HEK cells significantly reduced carbachol-induced mitochondrial Ca²⁺ uptake which was significantly restored by re-introduction of INF2-caax but not INF2-non-caax (**Figures 5E and 5F**). Interestingly, Store Operated Calcium Entry (SOCE) induced mitochondrial Ca^2+^ uptake was unperturbed in the INF2-KO cells suggesting that INF2 specifically altered ER to mitochondrial Ca^2+^ transfer (**Figure 5G**). Therefore, it was evident that while ER localization of INF2 was not required for agonist-induced IP3R2 activity, INF2-caax specifically influenced the ability of IP3R2 to mediate ERMC and downstream agonist-induced mitochondrial Ca²⁺ uptake. This further prompted us to test whether INF2 have IP3R-independent role in regulation of ERMC. PLA assays showed a significant reduction in the abundance of PLA dots but not PLA density upon INF2 depletion (KO) in IP3R-TKO HEK cells suggesting that INF2 might have a general role in mediating ERMC (**Figures 5H and 5I**). To dissect this further we evaluated the extent of ERMC in these cells from thin-section TEM. Interestingly, chronic INF2-depletion significantly reduced close ERMC (10-20 and 20-30 nm) while still preserving the tightest contacts (0-10 nm) in these cells (**Figures 5J and 5K**). Collectively these results further confirm that INF2 plays a pivotal role in mediating ERMC and downstream agonist-induced mitochondrial calcium uptake.

**Figure 5:**
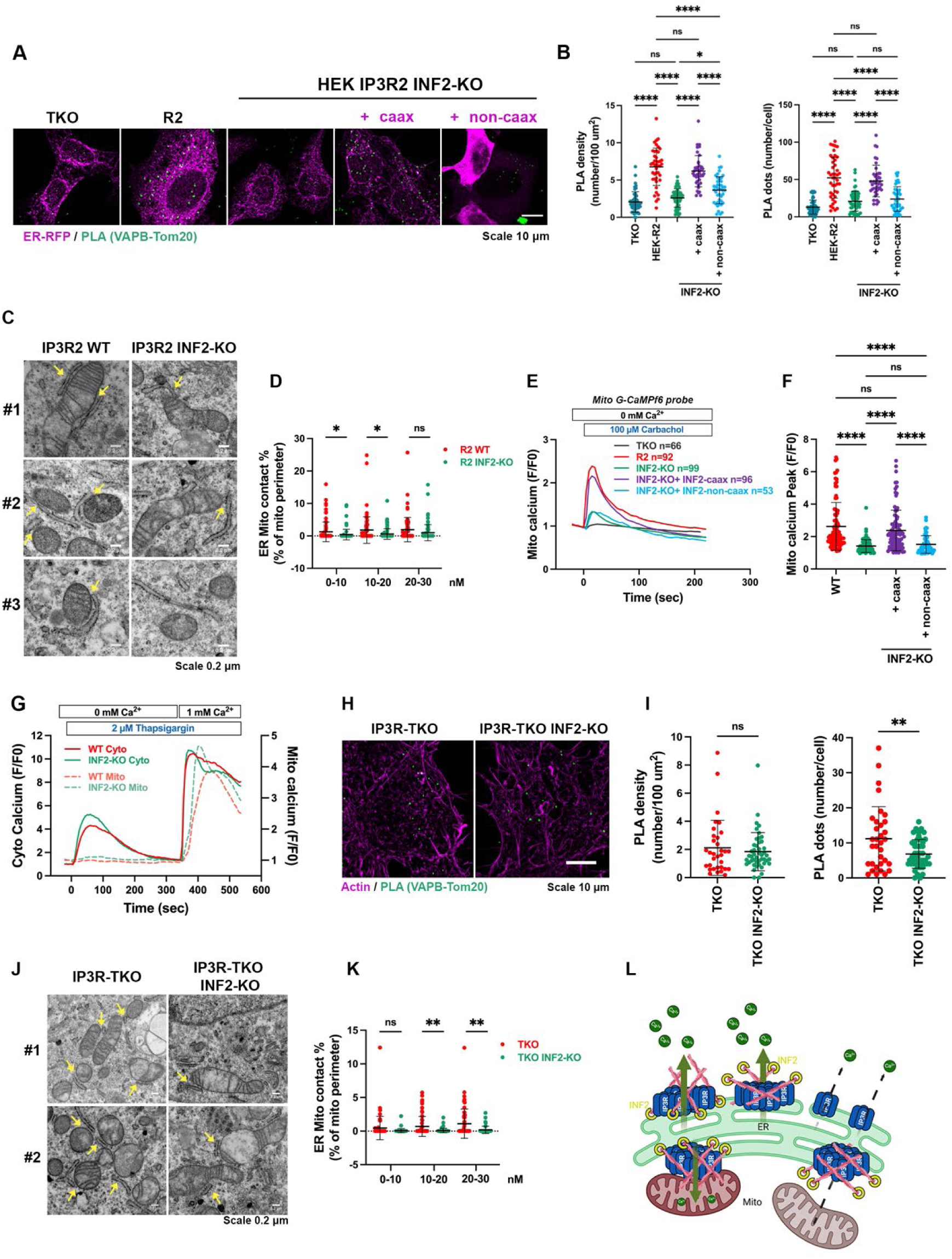
INF2-caax specifically regulates ER-Mito contacts and ER to mito calcium transfer. A: Representative confocal micrographs from PLA assay against VAP-B (ER) and Tom20 (mitochondria) to assess ERMC, in HEK IP3R-TKO, IP3R2 (R1/R3 DKO) WT / INF2-KO and INF2-KO cells rescued with INF2-caax or INF2-noncaax. TKO, WT and INF2-KO cells were transfected with ER-RFP (magenta) to label ER prior to PLA staining. The PLA dots of VAP-B/Tom20 proximity is shown in green. Scale: 10 µm. B: Quantification of PLA density expressed as number of dots per 100 µm2 (left), and PLA dots, expressed as number/cell (right). Data from 49 cells for TKO, 43 for WT, 59 for INF2-KO, 40 for INF2-KO + INF2-caax and 43 INF2-KO + INF2 noncaax from 3 independent experiments. Graphs show Mean ± S.D. One-way Anova used. **** p<0.0001, *** p=0.0004, * p=0.0370 (density); **** p<0.0001, * p=0.0367 (dots). C: Representative images from thin-section TEM in WT and INF2-KO HEK-IP3R2 (R1/R3 DKO) cells showing ER and mitochondria in the sections. The arrows represent close contact between ER and Mitochondria in the sections. Scale: 200 nm. D: Scatter plots showing the percentage of close ER-Mito contacts, expressed as % of mitochondrial perimeter, between 0-10, 10-20 and 20-30 nm. Data of 79 (WT) and 89 (INF2-KO) mitochondria, from 2 independent sets of images. Graphs show Mean ± S.D. unpaired Student’s T-test. * 0.0251 (WT/INF2 KO 0-10 nm) and * 0.0106 (WT/INF2 KO 10-20 nm). E: Traces of mitochondrial calcium transients measured in HEK IP3R2 (R1/R3 DKO) WT, INF2-KO and INF2-KO rescue (+caax or +noncaax) cells, transfected with Mito GCaMPf6 (Kd=0.2 µM). F: Quantification of maximal peak of mitochondrial Ca^2+^ uptake in C, expressed as fold change F/F0. Graphs show Mean ± S.D. Significant was measured using One-way Anova. Data from 66 (IP3R-TKO), 92 (IP3R2 expressing HEK), 99 (IP3R2 expressing INF2-KO), 96 (INF2-KO + INF2 caax) and 53 (INF2-KO + INF2-noncaax) across 3-4 independent experiments. Graphs show Mean ± S.D. Significant was measured using One-way ANOVA. **** p<0.0001. G: Traces of cytosolic (Cyto-R-GECO) and mitochondrial (MitoGCaMPf6) calcium transients measured simultaneously in WT and INF2-KO cells following 2 uM Thapsigargin treatments in calcium free ECM followed by re-introduction of 1 mM CaCl_2_. Traces shown are average value from 38 WT and INF2 KO cells across 2 independent experiments. H: Confocal micrographs from PLA against VAP-B (ER) and TOM20 (mitochondria) to assess ERMC, in HEK TKO and HEK TKO INF2-KO. Cells were transfected with ER-RFP (magenta) to label ER prior to PLA staining. The interaction of these proteins is shown in green dots. Scale: 10 µm. I: Quantification of PLA density expressed as number of dots per 100 µm2 (left), and PLA dots, expressed as number/cell (right). Data of 33 cells for TKO and 46 TKO INF2-KO, from 3 independent experiments. Graphs show Mean ± S.D. Significant was measured using unpaired Student’s t-test. Graphs show Mean ± S.D. One-way ANOVA used. ** p=0.0054. J: Representative images from thin-section TEM in IP3R-TKO HEK and IP3R-TKO-INF2-KO HEK cells showing ER and mitochondria in the sections. The arrows represent close contact between ER and Mitochondria in the sections. Scale: 200 nm. K: Scatter plots showing the percentage of close ER-Mito contacts, expressed as % of mitochondrial perimeter, between 0-10, 10-20 and 20-30 nm. Data of 60 (TKO) and 55 (TKO-INF2-KO) mitochondria, from 2 independent sets of images. Graphs show Mean ± S.D. unpaired Student’s T-test. ** 0.0034 (TKO/TKO-INF2 KO 10-20 nm) and ** 0.003 (TKO/TKO-INF2 KO 20-30 nm). L: Working model for INF2-dependent actin-mediated clustering, positioning and activity of IP3R in cells.

## Discussion

Collectively our data shows that INF2-mediated actin filaments play an integral role in agonist-induced IP3R-mediated ER calcium release. INF2-localization on the ER is dispensable for its role in IP3R activity. Our study further shows that depletion of INF2 significantly reduced IP3R clustering which have been shown to be important for regulation of IP3R. Further, both INF2 isoforms can significantly interact with all IP3R isoforms in an activity and actin filament independent manner, however, INF2 activity is imperative for IP3R cluster formation and agonist-mediated firing of IP3Rs. Finally, we show that the ER localization of INF2 influences proper positioning of IP3R2 close to mitochondria, mediating ER-Mitochondrial contacts and promoting ER to mitochondrial calcium transport. Overall, we show that in addition to previously described role of INF2 in mitochondrial division, it plays additional role upstream in the process by regulating (1) IP3R clustering; (2) IP3R-mediated ER calcium release and (3) IP3R positioning leading to ER-Mito contacts (**Figure 5L**).

IP3Rs in cells are shown to be arranged in clusters ^36,37^ and further have been classified into mobile and active immobile clusters ^36^. Ca^2+^ ^38^, an open transition state of IP3R ^39^, agonist-induction ^40^ and IP3 ^41^ have been shown to regulate clustering which is important for IP3R activity. It is, however, difficult to imagine what regulates the formation or lends stability to these clusters to allow Ca^2+^ release. Our results show that depletion of INF2 or its activity significantly reduces abundance of IP3R2 and IP3R3 clusters on ER signifying that actin filaments play an important role in regulating the clusters in the following possible ways: (1) INF2-mediated actin filaments might generate force needed for cluster formation; (2) INF2-mediated actin filaments might act as a scaffold required for cluster stability through either a force dependent or independent mechanism. In fact, actin filaments have been implicated in regulating functional clusters formation of Fcɣ receptors ^42^, CD44 on PM ^43^, influenza hemagglutinin ^44^ and RNA polymerase II in the nucleus ^45^. While INF2 can polymerize actin, extensive actin bundling which often prevents its cofilin-mediated depolymerization ^46^ might be required to aid in IP3R clustering which need to be studied further. Microtubules have been shown to positively regulate agonist-induced Ca^2+^ spike in epithelial cells ^47^ and a recent study have shown that depletion of microtubule (+) end binding protein EB3 significantly decreased IP3R3 cluster lifetime ^48^ raising speculations that actin and microtubule cytoskeleton might work in sync to regulate cluster assembly. At present it is difficult to tease out whether INF2-mediated actin filaments work upstream of EB3; however, since multiple formin proteins including INF2 have been shown to bind and modulate microtubules dynamics ^49–51^, it is possible that actin filaments might also allow for IP3R-EB3 binding to allow for greater stability of the IP3R clusters. Furthermore, transition from mobile to immobile clusters have been shown to be mediated by KRAP-induced tethering of IP3R clusters with actin filaments in the PM at sites of Store operated Ca²⁺ entry ^52^ which is likely to be downstream of INF2-mediated assembly of IP3R clusters. Overall, we think that actin filaments, possibly together with the microtubule network, might act as a scaffold to allow for IP3R cluster assembly and stability to facilitate agonist-induced ER Ca²⁺ release.

Through multiple approaches we show a direct binding of INF2 with IP3R isoforms in cells, however, the exact domains involved in this interaction remains to be worked out. Clearly the N-terminal DID region of INF2, and the actin polymerization activity is dispensable for its binding with IP3R (**Figure 3)** suggesting that FH1, FH2, DAD or the C-terminal region post DAD domain might be important for the interaction. In fact, we show that INF2-caax S1099A and S1100A mutants are significantly compromised for their ability to bind to IP3R2 further implicating that the C-terminal tail might be an important regulation hub for INF2. However, further analysis is required to (1) identify additional factors that might aid in INF2-IP3R interaction; (2) dissect out whether a double S1099/1100A mutant totally abrogates interaction; (3) whether the amino acid residues per se and not the phosphorylation is what determines the interaction. Unlike INF2, IP3R boasts of an exhaustive list of interactors ^53^ and through this study we add INF2 to the growing list. Clearly IP3 binding and pore-forming ability of IP3R does not influence its binding with INF2. An important molecule sitting at the speculative interface of this binding might be calmodulin which is known to bind IP3R in an inhibitory way ^54^ but has been shown to activate INF2 ^27–31^ in a Ca²⁺-dependent manner. It remains to be seen whether INF2-calmodulin binding allows sequestration of calmodulin from IP3R thereby further facilitating Ca²⁺ release.

In addition to mitochondrial fission, we show that INF2 regulates formation of ERMC. Interestingly our previous reports using U2-OS cells ^26^ showed an unchanged basal ERMC upon INF2-KO, however both in unstimulated HeLa and HEK-293 cells we see a drastic reduction of close ERMC upon INF2 depletion both at the level of light microscopy and TEM analysis (**Figure 5**). This might be due to a cell-to-cell variation or method of quantification employed in the two studies. We further show that INF2 depletion significantly dampened the ability of IP3R2 to facilitate ERMC (**Figure 5**) which is probably due to a reduction in IP3R cluster formation. While all IP3R isoforms have been shown to form clusters close to mitochondria ^23^, IP3R2 seemed to have a higher overall contacts with mitochondria ^23^. Our results further demonstrate that ER localized INF2-caax specifically regulated positioning of IP3R2 clusters close to mitochondria thereby facilitating not only the formation ERMC but also ER to mitochondrial Ca²⁺ transport. In fact, capture of IP3R clusters at ERMC allows it to stimulate local Ca²⁺ transfer to mitochondria boosting oxidative metabolism ^55^ implicating the importance of INF2-mediated ER bound actin filaments in the process. Since ER shaping proteins like REEP1 ^56^ and RTN1A ^57^ have been shown to regulate ERMC, INF2 mediated regulation of ERMC might be due to alteration of ER structure in INF2 depleted cells which is recovered by ER bound INF2-caax only. Does INF2 work only with IP3Rs to mediate ERMC? Most probably the answer is negative. A significant reduction of ERMC in INF2 depleted IP3R-TKO cells point to the fact that ER-bound INF2 might be a global regulator of ERMC in general. Therefore, in addition to positioning important ERMC molecules at the interface, INF2 on the ER might work with other members on the outer mitochondrial membrane like Spire1 ^58^ or Myosin-19 ^59^ to stabilize ERMC. Further work is necessary to elucidate the specific role of INF2 at ERMC and its interplay with various molecular determinants on ER that regulate ER-mitochondrial crosstalk.

## Materials and Methods

### Cell culture

Human cervical cancer cells (HeLa) were obtained from American Type Culture Collection (ATCC), and Human Embryonic Kidney (HEK) cells expressing specific IP3R isoforms were a kind gift of Dr. David Yule ^32^. Cells were grown in regular DMEM (Gibco 11-995-065) supplemented with 10% fetal bovine serum at 37°C with 5% CO2. Cell lines were tested every 3 months for mycoplasma contamination using Universal Mycoplasma detection kit (ATCC, 30-1012K) or MycoAlert Plus Mycoplasma Detection Ki (Lonza, LT07-701).

### Generation of INF2-KO cell lines

HeLa and IP3R isoform-specific HEK cells were genome-edited for deletion of INF2 gene using the CRISPR/Cas9 technology. CRISPR-gRNA for INF2 was as follows: 5’- CACCG CGGAGATACGTGCAACGCCG -3’ that was designed to target a specific genomic region in exon 1. Selected gRNAs were cloned into pSpCas9(BB)-2A-Puro (Addgene #48139), sequence-verified, and 1 µg of respective plasmids were transfected using Lipofectamine2000. Next day, transfected cells were replated into medium containing 2.5 µg/ml puromycin (Gibco #A1113802) and selected for 24 hours. After 24 hours puromycin containing media along with dead cells were replaced with normal DMEM and the adherent cells were allowed to become confluent. The cells were then diluted to ∼0.5 cells per well and seeded in a 96 well to obtain single cell derived clones. Single cell clones were expanded and screened for the absence of INF2 protein expression by western blotting using specific antibodies.

### Plasmids

Constructs for calcium transients: Cyto-R-Geco (Kd= 0.5 µM) and Cyto LAR-Geco (Kd=12 µM) was a gift from Y.M. Usachev (University of Iowa Carver College of Medicine, Iowa City, IA), ER GCaMP6 150 (Kd= 150 µM) was a kind gift from Timothy Ryan (Weil Cornell Medicine, NYC, NY), CMV-ER-LAR-GECO1 was a gift from Robert Campbell (Addgene plasmid # 61244), Mito GCaMPf6 (Kd=0.2 µM) was a kind gift from Gyorgy Hajnoczky (Thomas Jefferson University, Philadelphia, USA). mcherry-Ftractin, GFP-INF2-caax, GFP-INF2-non-caax, GFP-INF2-caaxC1270S, GFP-INF2-caax-FFC, GFP-INF2 I676A, GFP-INF2-caax-S1099A, GFP-INF2-caax-S1100A, eGFP-Tom20, ER-RFP and GFP-Sec61β were a kind gift from Henry Higgs (Geisel School of Medicine, Dartmouth). IP3R1 plasmids encoding the D2550A pore mutant and the R265Q ligand-binding domain mutant were from Suresh Joseph (Thomas Jefferson University) and have been described previously ^60^. YFP-PH-PLCδ plasmid was a kind gift from Tamas Balla (NIH, Bethesda). ERM-APEX2 was a gift from Alice Ting (Addgene plasmid # 79055). To generate Apex2-IP3R1 construct, a 747bp DNA fragment was synthesized by Twist Biosciences (San Francisco, CA) containing the Apex2 enzyme attached to a portion of the N-terminus of IP3R1 by a linker (GGSGGGSG). The fragment was synthesized with flanking EcoRI/KpnI sites which were used to insert the fragment into the full length IP3R1 plasmid.

For all experiments, the following amounts of DNA were transfected per 6-well (individually or combined for co-transfection): 250 ng of Cyto-R-GECO, Cyto-LAR-GECO, mito-GCaMPf6, YFP-PH-PLCδ, mcherry-Ftractin; 500 ng of ER GCaMP6 150, ER LAR-GECO, ER-RFP, GFP-Sec61β; 1000 ng of GFP-INF2-caax, GFP-INF2-non-caax, GFP-INF2-caax C1270S and GFP-INF2-caax FFC; 800 ng of IP3R1-WT, IP3R1-D2550A, IP3R1-R265Q

### Plasmid transfections

Cells were seeded at 4×10^5^ cells per well in a 35 mm dish at ∼16h before transfection. Plasmid transfections were made in OPTI-MEM media (Gibco, 31985062) containing 2.5 µl of Lipofectamine 2000 (Invitrogen, 11668) following the manufacturer’s protocol. After 4-6 h of transfection, cells were trypsinized and re-plated onto glass bottomed dishes (MatTek Corporation, P35G-1.5-14-C) at ∼1×10^5^ cells per well (for live cell imaging) or onto coverslips (25 mm) for fixed cell analysis. Cells were imaged or fixed ∼16–24 h after transfection.

### siRNA induced INF2 knockdown

Cells were seeded at 1×10^5^ cells per well in a 35 mm dish ∼ 16 h before transfection. 63 pg of siRNA was added to OPTI-MEM (Gibco, 31985062) with 2 µl RNAimax (Invitrogen, 13778100), incubated at RT for 20 minutes and added to cells contained in 1 ml of pre-warmed DMEM media. 72-96 hours later cells were transfected with respective reporter constructs and prepared for live/fixed cell analysis. INF2-siRNA oligo: 5’-GGAUCAACCUGGAGAUCAUCCGC-3′

### Drug treatments

To evaluate the agonist-induced calcium release from ER histamine (H7125; Sigma-Aldrich) or Carbachol (thermo scientific 51-83-2) were used at different concentrations. Agonist was perfused manually at the fifth frame (∼25 sec) during the imaging and not replaced during the entire protocol (5 min). To evaluate basal cytosolic calcium levels, cells were loaded with 2 µM FURA-2 AM (Thermo Fisher Scientific, F1221). To evaluate the ER calcium stores, 2 µM of Thapsigargin (MilliporeSigma, 5860051MG) was added at the thirtieth frame (∼2 min) during the imaging and continued for another 5 min. For the Latrunculin A (Thermo Fisher Scientific, L12370) treatment, cells were pre-treated in DMEM with 0.5 µM of drug during 15 min, washed with PBS1X, and then the samples were collected following the IP protocol.

### Immunofluorescence

For IP3R cluster, ER structure, HeLa and HEK cells were subjected to immunofluorescence staining. Cells (transfected or un-transfected) were plated onto coverslips ∼16 h prior to fixation and staining. Cells were then fixed with chilled Methanol (for IP3Rs clusters, for 10 min at -20C) or with 4% PFA prewarmed (rest of IF, for 20 min at room temperature). In case of PFA fixation, cells were washed with PBS, permeabilized with 0.1% Triton X-100 for 1 min and washed with PBS three times. In all conditions cells were blocked with 10% FBS in PBS for ∼60 min. Primary antibody was diluted in 1% FBS/PBS and coverslips were incubated on a drop of antibody solution on parafilm in a wet chamber for 1 h. Cells were then washed 6 times with 1X PBS and incubated with appropriate secondary antibody (prepared in 1% FBS/PBS) for 1 hour. Coverslips were again washed 6 times with 1X PBS and mounted on glass slides using ProLong Gold antifade mounting media (Invitrogen #P36930).

#### Primary antibodies

anti-INF2 antibody (Proteintech, 20466-1-AP, used at 1:250)

anti-IP3R2 (A-5) antibody (Santa Cruz Biotechnologies, sc-398434, used at 1:100)

anti-IP3R3 antibody (BD Biosciences, 610312, used at 1:100)

anti-Tom 20 (Abcam ab78547, used at 1:400)

#### Secondary antibodies

Goat anti-Mouse IgG (H+L) Cross-Adsorbed Secondary Antibody, Alexa Fluor™ 647 (Invitrogen #A21235) used at 1:1000; Goat anti-Rabbit IgG (H+L) Cross-Adsorbed Secondary Antibody, Alexa Fluor™ 488 (Invitrogen #A11008) used at 1:1000

### Proximity Ligation Assay (PLA)

PLA (Duolink™ PLA technology, *In Situ* Starter Kit Mouse/Rabbit, MilliporeSigma, cat# DUO92101) was used to evaluate the interaction between INF2/IP3Rs or VAPB/Tom20. Briefly, cells were seeded onto 12 mm coverslips for 16 hours and then were fixed with chilled methanol 10 min/-20C (for INF2/IP3Rs), or with prewarmed 4% PFA for 20 min at RT (to VAP-B/Tom20). Cells were washed with PBS (X 3 times), and in case of PFA fixation, cells were permeabilized using 0.1% Triton-X-100 for 1 min. The cells were then incubated with Duolink blocking buffer for 1 hour and incubated with primary antibodies (diluted in Duolink antibody diluent) for 2 hours at RT. Cells were then washed with 5% BSA (15 min x 3) and incubated with Duolink secondary antibodies for 1 hr at 37C. Cells were then washed with Duolink wash buffer A (5 min x 3) and incubated with Duolink ligase for 30 min at 37C. Cells were further washed with Duolink wash buffer A (2 min x 3) and incubated with Duolink polymerase for 90 min at 37 C. The cells were finally washed with Duolink wash buffer B (10 min x 3) and mounted on glass slides using prolong gold antifade reagent. Images were acquired within 24-48 hours. For multiplexed PLA and IP3R3 staining, Alexafluor-647 coupled secondary antibody was also added at 1:2000 during the secondary antibody incubations to visualize IP3R3 along with the PLA dots. For VAP-B/Tom20 PLA staining, the Alexa fluor 488 Phalloidin was added in the blocking buffer to mark actin filaments.

#### Antibodies and stains used

anti-INF2 (Proteintech, 20466-1-AP, used at 1:250)

anti-IP3R2 (A-5) antibody (Santa Cruz Biotechnologies, sc-398434, used at 1:100)

anti-IP3R3 antibody (BD Biosciences, 610312, used at 1:100)

anti-VAP-B (Proteintech 66191-1-Ig used at 1:400)

and anti-Tom20 (Abcam ab78547, used at 1:400)

Alexa fluor 488 Phalloidin (Invitrogen, A12379, used at 1:500)

### Microscopy

#### Agonist-induced calcium dynamics

The images were acquired using a Nikon A1 on a Nikon Ti-E base and equipped with an iXon Ultra 888 EMCCD camera. A solid-state 405 smart diode 100-mW laser, solid-state 488 OPSL smart laser 50-mW laser, solid-state 560 OPSL smart laser 50-mW laser, and solid-state 637 OPSL smart laser 140-mW laser were used (objective: 20X, 1.45 NA CFI Plan Apo; Nikon). Images were acquired using Nikon Elements v5.02. Images from Fig. 1, 2, 4I, 4J, 5C, 5D, and Fig. S1B, S1H, S1I, S3, S5B and S5C were acquired using this system and quantified using Image J.

#### Basal calcium estimation

Epifluorescence Ca^2+^ imaging for *SI Appendix*, Fig. S1C, S1D,S1E, S2C, S2D and S2E, were acquired using the EM-CCD camera (Hamamatsu) assembled to an Olympus IX81 microscope with Lambda DG4 light source operated by Metamorph (Molecular Devices). Time lapse images were acquired using the UV-optimized Olympus UAPO/340 ×40/1.35 N.A. oil immersion objective every 4 s.

#### Fixed samples

Was performed on LSM 880 equipped with 63×/1.4 NA plan Apochromat oil objective using the Airyscan detectors (Carl Zeiss Microscopy). The Airyscan uses a 32-channel array of GaAsP detectors configured as 0.2 airy units per channel to collect data that is subsequently processed using Zen2 software. Cells were imaged with the 405-nm laser and 450/30 filter for BFP, 488 nm and 525/30 for GFP, 561 nm and 595/25 for RFP and 640 nm laser and 650 LP for Far-red. Images for Fig. 3A, 4C, 4E, 4G, 5A 5E, Fig. S1B, S4A, S5E, S5F, S5H, S5J and S6A, were acquired using this system and quantified using Image J.

### Calcium measurements

Cells expressing respective calcium sensors either alone or with INF2-rescue plasmids were washed twice with PBS1X and perfused with 0.25 % BSA ECM buffer, calcium-free (ECM: 121 mM NaCl, 5 mM NaHCO_3_, 10 mM Na-HEPES, 4.6 mM KCl, 1.2 mM KH_2_PO_4_, 1.2 mM MgSO_4_, 10 mM glucose, pH7.4) and transferred to the microscope equipped with a heated (37C) stage-top. Cells were recorded for 25 seconds at 5s/frame, stimulated with agonists and imaged for another 5 minutes at 5s/frame. Images were quantified using ImageJ where values were expressed as fold change F/F0 where F0 is the mean of resting fluorescence intensity before stimulus addition. For rapid kinetics, cells expressing Cyto-R-Geco were recorded at 1frame/sec for 4 minutes.

To determine the basal cytosolic and ER Ca^2+^ levels, cells were loaded with 2 µM Fura-2 AM in the presence of 0.003% Pluronic acid F-127 and 100 μM sulfinpyrazone for 10 min at room temperature, washed with fresh ECM containing 0.25% BSA, calcium-free, and transferred to the microscope stage (37C). Fluorescence was recorded at 340 and 380 nm excitation, using dual-band dichroic and emission filters at 4s/frame. To evaluate the ER stores, 2 µM of Thapsigargin was added after 2 min of initial recording and further imaged for another 8 min. Values were expressed as F340/F380 ratio.

### Actin burst

To measure actin burst, U2OS-INF2-KO cells were co-transfected with GFP-INF2 constructs the day before of imaging. After ∼16–24 h of transfections, cells were stimulated with 4 uM ionomycin for 45 seconds, fixed with 4% pre-warmed PFA for 20 min at RT, washed with PBS (X3), permeabilized with 0.1% Triton X-100 for 1 min at RT, washed with PBS (X3) and stained with phalloidin-647 for 30 minutes at RT. The cells were then washed with PBS and mounted on slides to be imaged.

### Lysate preparation and Western Blot

To evaluate protein abundance (except IP3R), cells from a 35mm dish were lysed using 1% SDS, collected and heated at 95C for 5 min. To evaluate abundance of IP3R, cells were lysed in RIPA buffer (reconstituted with 1X Protease/Phosphatase inhibitors) and saved at -80C without boiling. To run them on SDS-PAGE, samples were mixed with 2xDB (50mM Tris-HCl, pH 6.8, 2mM EDTA, 20% glycerol, 0.8% SDS, 0.02% Bromophenol Blue, 1M NaCl, 4M urea) and separated in a 4-20% precast gradient gels (all proteins except IP3R), or custom-made 5% Tris-glycine gels (in case of IP3Rs), transferred to a polyvinylidene fluoride membrane (EMD Millipore, IPFL00010). The membrane was blocked with TBS-T (20 mM Tris-HCl, pH 7.6, 136 mM NaCl, 0.1% Tween20) containing 3% BSA (VWR Life Science, VWRV0332) for 1h, then incubated with primary antibody solution at 4°C overnight or 2 hours at RT. After washing with TBS-T, the membrane was incubated with requisite secondary antibody for 1h at room-temperature and developed on a Chemi-doc system using appropriate reagents.

#### Primary Antibodies used

anti-INF2 antibody (Proteintech, 20466-1-AP, used at 1:1000)

anti-β-Actin antibody (CST, #3700S, used at 1:5000; SIGMA, A2228 used at 1:10000)

anti-IP3R1 (D53A5) antibody (CST, 8568S, used at 1:1000)

anti-IP3R2 (A-5) antibody (Santa Cruz Biotechnologies, sc-398434, used at 1:500)

anti-IP3R3 antibody (BD Biosciences, 610312, used at 1:2000)

anti-VDAC (SA93-03) antibody (Invitrogen, MA5-41088, used at 1:1000)

anti-VAP-B (Proteintech 66191-1-Ig used at 1:2000 at RT/ 2hr)

anti-Tom20 (Abcam ab78547, used at 1:1000)

anti-PTPIP51 (Proteintech 20641-1-AP used at 1:1500 at RT/ 2 hr)

anti-IRBIT (Cell Signaling Technology #94248 used at 1:1000)

anti-α-actinin (Cell Signaling Technology #6487 used at 1:1000)

#### Secondary Antibodies used

anti-Rabbit-HRP antibody (Jackson Immuno Research, 111-035-144, used at 1:10000)

anti-Mouse-HRP (Jackson Immuno Research, 715-035-150, used at 1:5000)

anti-rabbit IgG (H+L) antibody conjugated to IRDye 800CW (LICORBio, 926-32211, used at 1:10000)

anti-Mouse IgG (H+L) antibody conjugated to IRDye 680LT (LICORBio, 926-68020, used at 1:10000)

### Immunoprecipitation using GFP-Trap agarose beads

INF2-IP3R2 interactions was measured using the GFP-Trap agarose beads (ChromoTek GFP-Trap® Agarose). Briefly, IP3R2 (IP3R1/3 DKO)-INF2-KO HEK cells were seeded into a 10 cm dish and were transfected with GFP-tagged-INF2-constructs. After 6h of transfection, media was replaced with fresh DMEM. Next day, media was removed, washed with PBS and lysates was prepare using immunoprecipitation buffer (IPB) (IPB: 140 mM KCl, 50 mM Tris-HCl (pH 7.4), 2 mM MgCl_2_) with protease and phosphatase inhibitors containing 1% Triton-X100, Lat A (2 µM) and Phalloidin (10 µM). Cells were lysed on a rotator for 1 h at 4C, then centrifugated for 15 min at 15000 rpm at 4C to collect the supernatant. ∼10% of the volume was used for the *Input*, and the rest was added to the conditioned GFP-trap-beads. After 2 h mixing on a rotator at 4C, the beads were collected via centrifugation, washed with IPB (X 3) and reconstituted in sample buffer, heated at 95C for 5min, and evaluated through SDS-PAGE.

For the pull-down with Lat A, transfected cells were pre-treated with 0.5 µM of Latrunculin A for 15 min at 37C and then processed as described above.

For pull-down using IP3R1 constructs, IP3-TKO-INF2-KO HEK cells transiently expressing IP3R1 constructs and GFP-INF2-caax were processed as above.

### APEX2-labelling for IP3R1

Proximity biotinylation was carried out using a variation of published protocol ^61^. Briefly, triplicate 100mm dishes of IP3R TKO cells were transfected with 10ug each of APEX2-IP3R1 or ERM-Apex2. After 48h, the cells were preincubated with 0.5mM Biotin phenol for 40min and then treated for 1min with 1mM H_2_O_2_. The biotinylation reaction was then quenched by washing in PBS containing 1mM of Trolox, sodium ascorbate and sodium azide. Cells were lysed in a buffer containing 150mM NaCl, 20mM Tris Hepes (pH 7.8), 0.5mM EDTA, 1% Triton X-100, 0.1% SDS, protease inhibitors and biotinylation quenching reagents. Biotinylated proteins in the lysates were recovered by incubation for 4h with streptavidin agarose beads. The beads were washed as described in Figure 3G and subjected to overnight digestion with trypsin. Samples were processed for LC-MS/MS at the Wistar proteomic facility.

### Transmission Electron Microscopy

HEK cells (**Figures *5C, D, I, J***) for TEM analysis were fixed using 2% glutaraldehyde, then with 1% OsO_4_ and stained with 0.5% uranyl acetate, pelleted in 2% agarose (Sigma–Aldrich, Type IX ultralow gelling temperature), dehydrated in a dilution series of acetone/ water, and embedded in Spurr’s resin (Electron Microscopy Sciences). The sections were examined with a JEOL JEM1010 TEM fitted with a side-mounted AMT XR- 50 5Mpx CCD camera or an FEI Tecnai 12 TEM fitted with a bottom-mounted AMT XR-111 10.5 Mpx CCD camera. Images were taken with or ×3200–4400 (FEI) direct magnification. HeLa WT and INF2-KO cells (**Figures S6C-S6E**) for electron microscopic examination were trypsinized, washed twice with PBS and fixed with 2.5% glutaraldehyde, 2.0% paraformaldehyde in 0.1M sodium cacodylate buffer, pH7.4, overnight at 4C in a rotator. After subsequent buffer washes, the samples were post-fixed in 2.0% osmium tetroxide for 1 hour at room temperature and rinsed in dH2O prior to *en bloc* staining with 2% uranyl acetate. After dehydration through a graded ethanol series, the cells were infiltrated and embedded in EMbed-812 (Electron Microscopy Sciences, Fort Washington, PA). Thin sections were stained with uranyl acetate and lead citrate and examined with a JEOL 1010 electron microscope fitted with a Hamamatsu digital camera and AMT Advantage NanoSprint500 software. Interfaces between ER and mitochondria were analyzed using a custom ImageJ script: https://sites.imagej.net/MitoCare/ as described elsewhere ^62^

### Image analysis and quantification

Image J (NIH) was used to analyze and process all images

#### Quantification of calcium transients

Cells transfected with the indicated calcium probes as described above were plated on MatTek dishes for live-cell imaging. Cells were imaged using a 20X water objective on a heated stage at a single confocal slice in the medial region, approximately 2 μm above the basal surface, to avoid stress fibers. Mean fluorescence intensity was calculated for each cell using the ImageJ plug-in “Time Series Analyzer V3.” after background subtraction (taken from extra-cellular 2-3 ROI in the field). Fluorescence values for each time point after drug treatment were normalized to the mean baseline fluorescence (first five frames) and plotted as intensity fold change over time in seconds. Average fluorescence intensities for mitochondrial calcium and ER calcium were determined by outlining the respective organelle networks. Data were plotted as intensity–time curves each data point using Microsoft Excel.

Cells loaded with FURA-2AM to determine the basal and ER Ca^2+^ stores, were analyzed using the custom-made Spectralyzer (imaging software). A masking for each cell was created and average intensities for F340 and F380 were collected and plotted as a ratio (F340/F380). An individual extra-cellular ROI was evaluated for both wavelengths to determine the background which was factored in the quantification.

#### Quantification of IP3R Clusters

z-projected images were background-subtracted using “rolling ball radius” plugin of Image J and threshold corrected to preserve the brightest cluster of IP3R. Similar parameters were used for all conditions imaged. The thresholded IP3R image was converted to 8 bit and binarized. “Analyze Particles” plugin was used to record the total number of clusters (as dots) and presented as number per 100 µm^2^ of cellular area. All images collected were quantified.

#### Quantification of PLA assays

z- projected images were background-subtracted using “rolling ball radius” plugin of Image J with a value of 20.0 and phalloidin (actin) channel was used to get the cellular area. Channel containing the PLA dots were converted to 8-bit and binarized. “Analyze particles” plugin was used to record total number of dots and presented as number of dots per 100 um^2^ of cellular area. All images collected were quantified.

### Statistical analysis

All statical analysis and p-values were determined using GraphPad Prism 10. To determine *p*-values, an unpaired Student’s *t*-test was performed between two groups of data, comparing full datasets as stated in the Figure legends. To compare across more than two conditions, multiple comparisons (unpaired) and one-way or two-ways ANOVA were performed in GraphPad Prism 10. Data are expressed as Mean±SD.

## Acknowledgments

We thank David Yule (University of Rochester) for kindly providing the IP3R expressing HEK cell lines, Tim Ryan (Weill Cornell Medicine) for sharing the ER-GCaMP150 calcium sensor, David Weaver for his expertise in imaging, Gyorgy Hajnoczky (Thomas Jefferson University), Henry Higgs (Geisel School of Medicine at Dartmouth) and Frieda Kage (Geisel School of Medicine at Dartmouth) for their critical advice and reagents, and to Pit Fin for his ubiquitous presence behind the scenes. We would also like to thank Zuzana Nichtova and Nina Petrova from Mitocare TEM Core facility and Biao Zuo and Inna Martynyuk from University of Pennsylvania - University of Pennsylvania Perelman School of Medicine Electron Microscopy Resource Lab Core Facility, RRID:SCR_022375 for their support in TEM studies. This work was supported by NIH R35 GM150811-3 (R.C)

## Author Contributions

M.R.Z., A.G., S.K.J., and R.C. designed research; M.R.Z., A.G., S.K.J., and R.C. performed research; A.G., and S.K.J. contributed new reagents/analytic tools; M.R.Z., A.G., S.K.J., and R.C. analyzed data; R.C. project supervision, funding acquisition and oversight; and M.R.Z., and R.C. wrote the paper

## Supplementary figures

**Figure S1:**
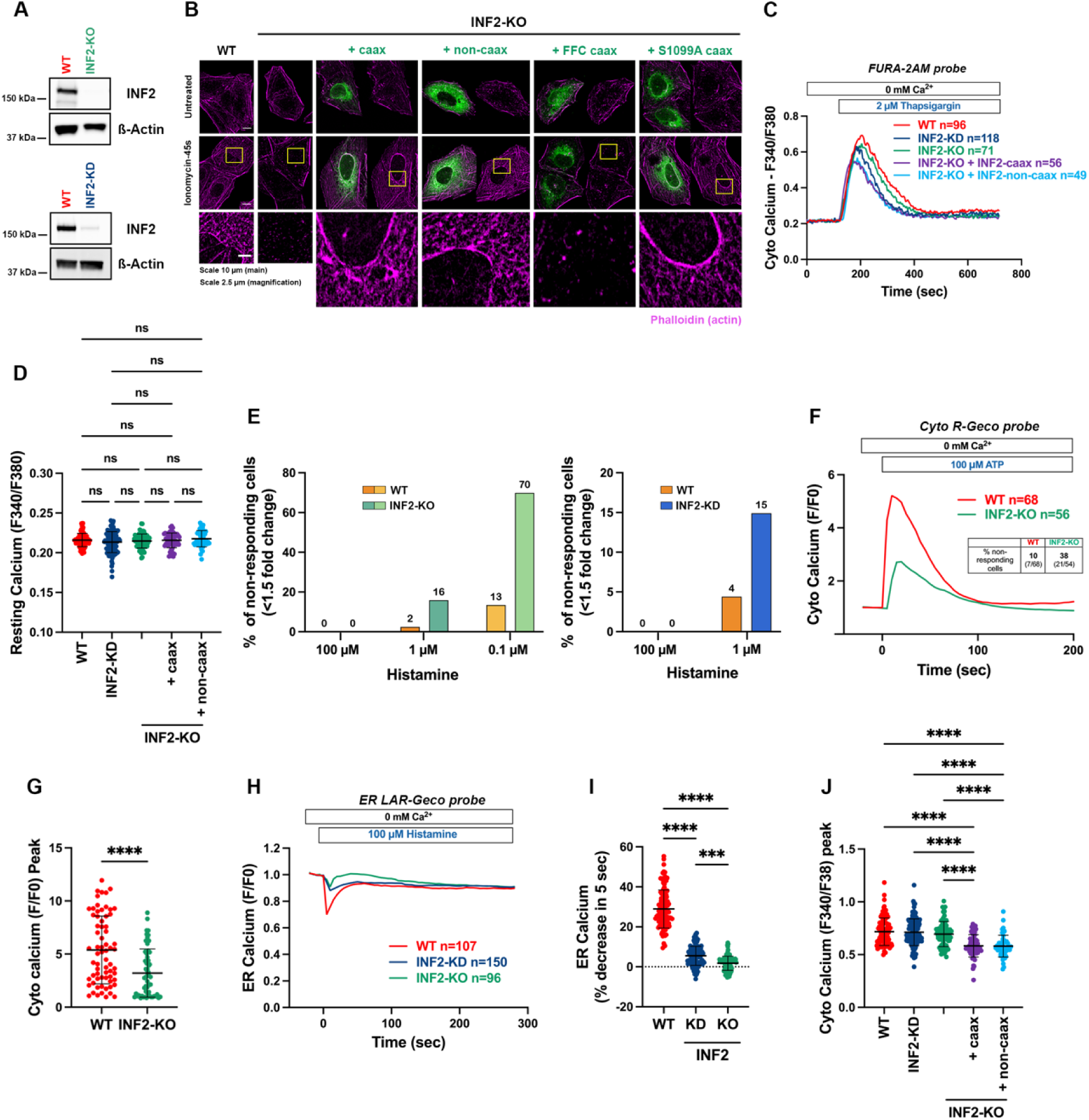
Depletion of INF2 reduces IP3R-mediated calcium release. A: Western blot to evaluate INF2 expression in WT, INF2-KO and INF2 KD in HeLa cells, β-actin used as loading control. B: Fixed-cell image of actin filaments (Phalloidin) in WT and INF2-KO U2OS cells either un-transfected or re-expressing INF2 constructs (WT-caax, WT-noncaax, FFC-caax and INF2 S1099A) before and after 45 seconds of stimulation with 4 µM ionomycin. Inset of the nucleo-cytoplasmic regions displaying the cytoplasmic actin burst is shown. Scale: 10 µm (Main) and 2.5 µm (inset) C: Cytosolic Ca2+ transients in HeLa WT, INF2-KD, INF2-KO and INF2-KO rescues cells, loaded with FURA-2AM, and stimulated with 2 µM of Thapsigargin to evaluate the ER stores. Cyto Ca2+ is expressed as F340/380. D: Resting calcium, F340/F380 determined as the average of the first 2 minutes of Cyto Ca2+, before the Thapsigargin addition. Data from 96 (WT), 118 (INF2-KD), 71 (INF2-KO), 56 (INF2-KO + INF2-caax) and 49 (INF2-KO + INF2-noncaax) cells across 3-4 independent experiments. Data shown as Mean ±S.D. One-way ANOVA used. **** p<0.0001 E: Quantification showing percentage of non-responding cells (fold change < 1.5) across all the conditions and histamine stimulations tested in Fig 1A-D. F: Traces of cytoplasmic calcium transients in HeLa WT and INF2-KO cells transfected with Cyto-R-GECO probe (Kd=0.5µM) and stimulated with 100 μM ATP. G: Quantification of maximal peak of cytosolic Ca^2+^ release in S1F after ATP treatment. Data from 68 (WT) and 56 (INF2-KO) HeLa cells across 2 independent experiments. Data shown as Mean ±S.D. One-way ANOVA used. **** p<0.0001 H: Traces of ER calcium transients in HeLa WT, INF2-KD and INF2-KO transfected with ER-LAR-GECO probe (Kd=24µM) and stimulated with 100 μM histamine. I: Graph quantifying acute change in ER calcium release upon the Histamine treatment, expressed as % of depletion at 5 sec. Data from 107 (WT), 150 (INF2-KD) and 96 (INF2-KO) HeLa cells across 3 independent experiments. Graphs show Mean ± S.D. One-way ANOVA used. **** p<0.0001, *** p=0.0001. J: Quantification of maximal peak of cytosolic Ca^2+^ release in S1C after Thapsigargin addition, expressed as F340/F380. Data from 96 (WT), 118 (INF2-KD), 71 (INF2-KO), 56 (INF2-KO + INF2-caax) and 49 (INF2-KO + INF2-noncaax) cells across 3-4 independent experiments. Data shown as Mean ±S.D. One-way ANOVA used. **** p<0.0001

**Figure S2:**
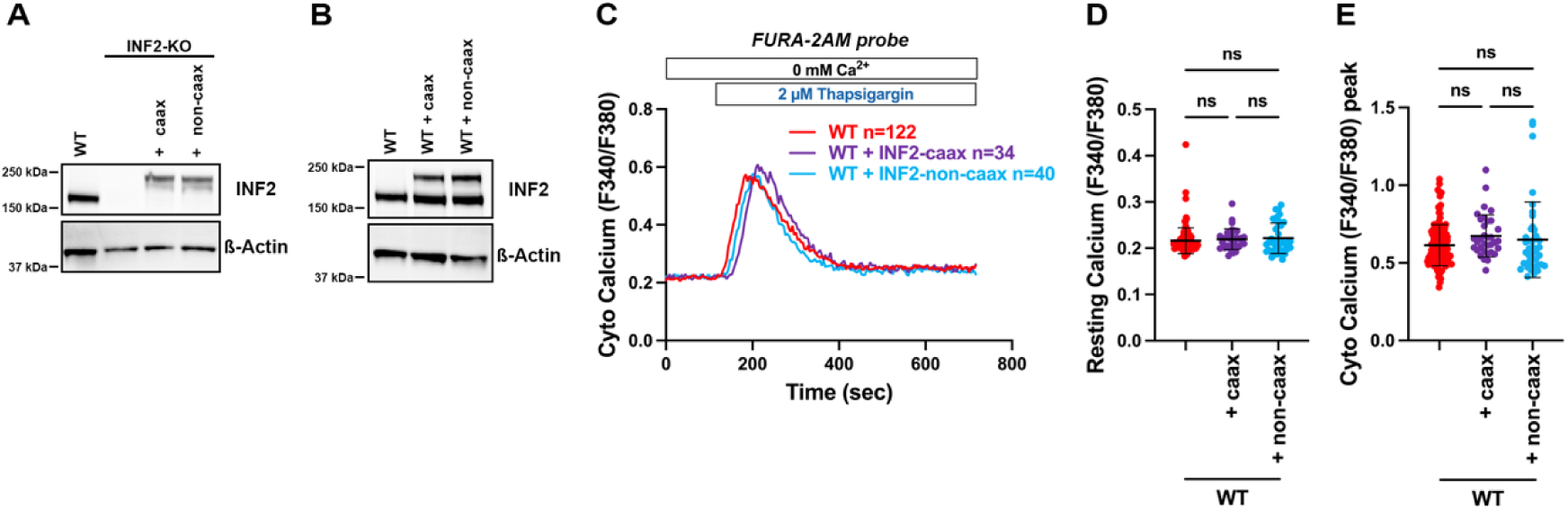
ER localization of INF2 is dispensable for IP3R-mediated calcium release. A: Western blot showing the INF2 expression in WT and INF2-KO HeLa cells transfected with INF2-caax and INF2-non-caax INF2-KO. β-actin used as loading control. B: Western blot showing the INF2 over-expression (INF2-caax and INF2-noncaax) in HeLA-WT cells. β-actin used as loading control. C. Cytosolic Ca2+ transients in HeLa WT and INF2 overexpressed cells, loaded with FURA-2AM, and stimulated with 2 µM of Thapsigargin to evaluate the ER stores. Cyto Ca2+ average was expressed as F340/F380. D. Resting calcium, F340/F380 determined as the average of the first 2 minutes of Cyto Ca^2+^ from C, before the Thapsigargin addition. Data from 122 (WT), 34 (WT + INF2-caax) and 40 (WT + INF2-noncaax) cells across 3 independent experiments. Graphs shown as Mean ± S.D. One-way ANOVA used. E. Quantification of maximal peak of cytosolic Ca2+ release in C after Thapsigargin addition, expressed as F340/F380. Data from 122 (WT), 34 (WT + INF2-caax) and 40 (WT + INF2-noncaax) cells across 3 independent experiments. Graphs shown as Mean ± S.D. One-way ANOVA used.

**Figure S3.**
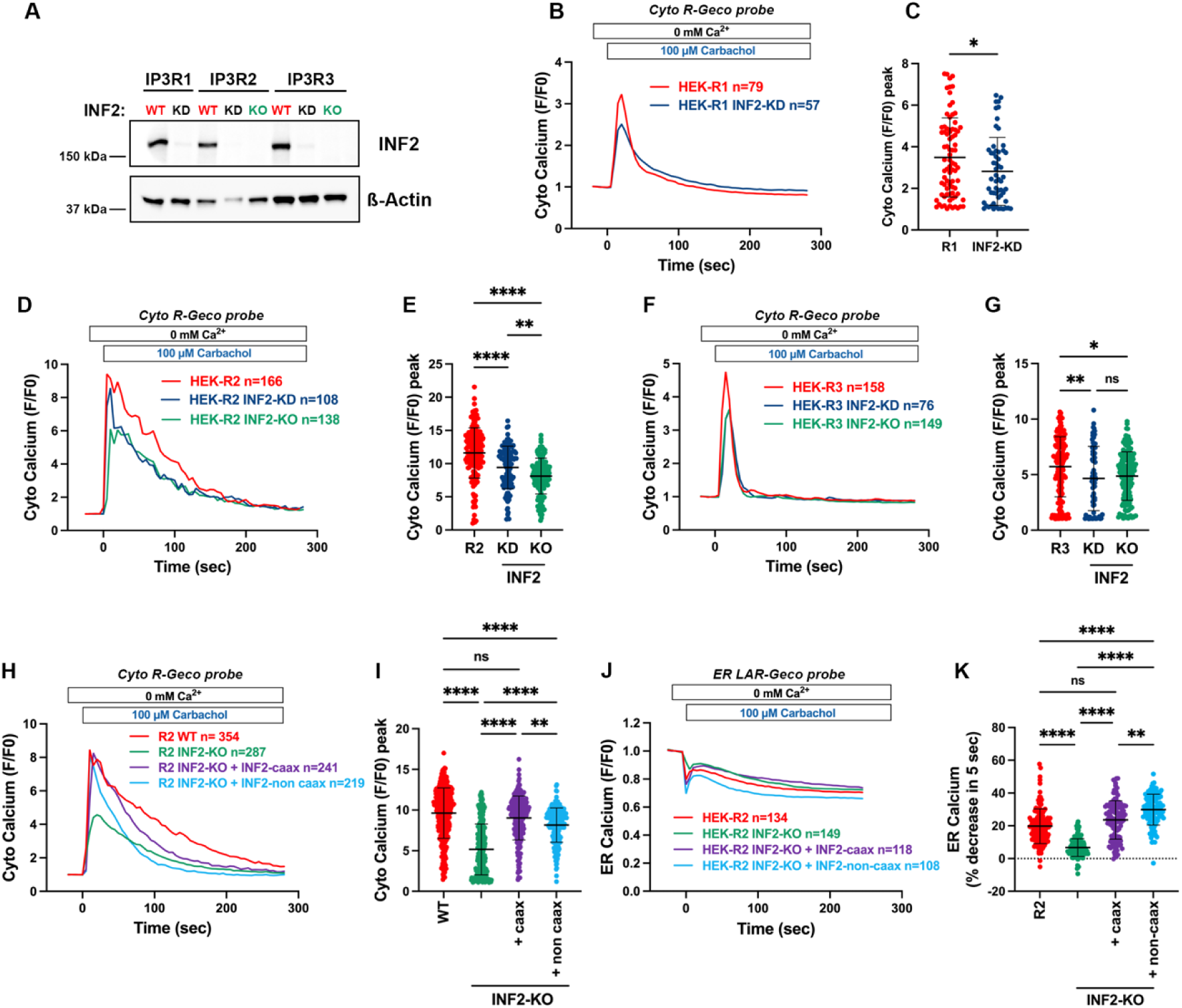
Depletion of INF2 affects the activity of all IP3R isoforms. A. Western blot showing the expression of INF2 in WT, INF2-KD OR INF2-KO done in HEK IP3R1 (R1) (R2/R3 DKO), IP3R2 (R2) (R1/R3 DKO) and IP3R3 (R3) R1/R2 DKO) as shown. Β-actin used as loading control. B: Cytosolic calcium transients in IP3R1 expressing HEK cells (WT and INF2 KD) transfected with Cyto-R-GECO probe and stimulated by Carbachol (100 µM). C: Quantification of maximal peak of cytosolic Ca^2+^ transients expressed as F/F0 in WT and INF2-KD HEK IP3R1 (R2/R3 DKO) cells. Data from 79 (WT) and 57 (INF2-KD) cells across 3 independent experiments. Graphs shown as Mean ±S.D. Students T-test used. * p= 0.0351. D: Cytosolic calcium transients in IP3R2 expressing HEK cells (WT, INF2 KD and INF2-KO) transfected with Cyto-R-GECO probe and stimulated by Carbachol (100 µM). E: Quantification of maximal peak of cytosolic Ca2+ transients expressed as F/F0 in WT and INF2-KD HEK IP3R2 (R1/R3 DKO) cells. Data from 166 (WT), 108 (INF2-KD) and 138 (INF2-KO) cells across 3 independent experiments. Graphs shown as Mean ±S.D. One-way ANOVA used. **** p<0.0001, ** p= 0.0038 F: Cytosolic calcium transients in IP3R3 expressing HEK cells (WT, INF2 KD and INF2-KO) transfected with Cyto-R-GECO probe and stimulated by Carbachol (100 µM). G: Quantification of maximal peak of cytosolic Ca^2+^ transients expressed as F/F0 in WT and INF2-KD HEK IP3R3 (R1/R2 DKO) cells. Data from 156 (WT), 76 (INF2-KD) and 149 (INF2-KO) cells across 3 independent experiments. Graphs shown as Mean ±S.D. One-way ANOVA used. ** p= 0.0097, * p=0.0110. H: Cytosolic calcium transients in WT and INF2 depleted IP3R2 expressing HEK cells rescued with INF2-caax or INF2-noncaax. The cells were transfected with Cyto-R-GECO probe and stimulated by Carbachol (100 µM). I: Quantification of maximal peak of cytosolic Ca^2+^ transients expressed as F/F0 from data set in S3H. Data from 354 (R2-WT cells), 287 (INF2-KO), 241 (INF2-KO + INF2-caax) and 219 (INF2-KO + INF2-noncaax) cells across 2-3 independent experiments. Graphs shown as Mean ±S.D. One-way ANOVA used. **** p<0.0001; ** p=0.0014 J: ER calcium transients in WT and INF2 depleted IP3R2 expressing HEK cells rescued with INF2-caax or INF2-noncaax. The cells were transfected with ER-LAR-GECO probe and stimulated by Carbachol (100 µM). K: Quantification of ER calcium release expressed as % of decrease in 5 sec for dataset in Fig S3J. Data from 134 (WT), 149 (INF2-KO), 118 (INF2-KO + INF2-caax) and 108 (INF2-KO + INF2-noncaax) cells across 3 independent experiments. Graphs shown as Mean ±S.D. One-way ANOVA used. **** p<0.0001, ** p=0.0012.

**Figure S4.**
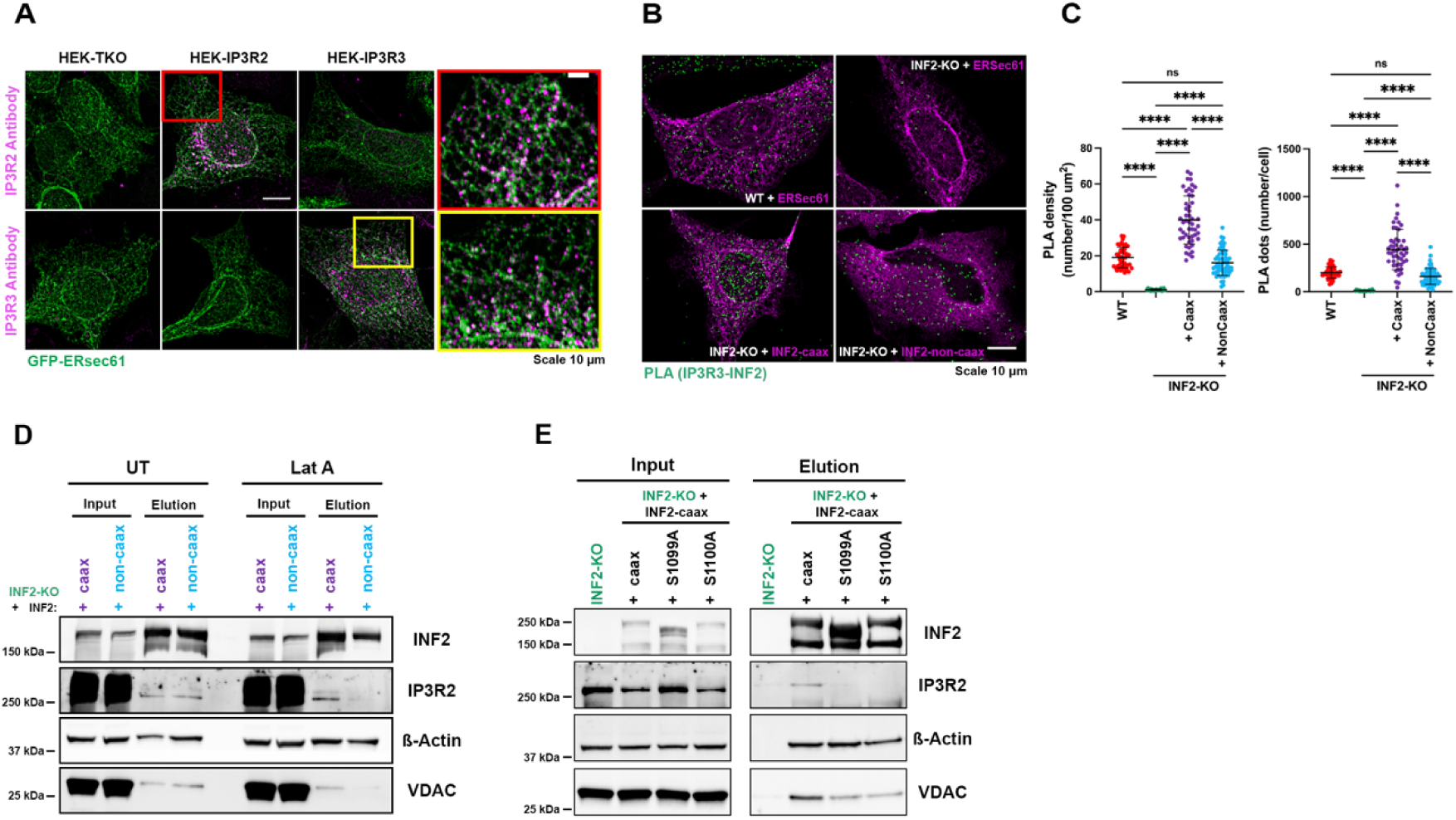
INF2 interacts with IP3R. A: Representative confocal micrograph of IP3R-TKO, IP3R2 (IP3R1/IP3R3 DKO) and IP3R2 (IP3R1/IP3R2 DKO) HEK cells transfected with ER-Sec61β (green), fixed and stained with IP3R2 and IP3R3 antibodies (magenta) as marked. Inset (Red) shows IP3R2 staining in IP3R2 expressing HEK cells and inset (cyan) shows IP3R3 staining in IP3R3 expressing HEK cells. B: Representative confocal micrographs from Proximity Ligation Assay (PLA) against IP3R3 and INF2 in HeLa WT, INF2-KO and INF2-KO rescue cells. WT and INF2-KO cells were transfected with GFP-ERSec61β to label ER (magenta) prior to PLA staining. The interaction of IP3R3/INF2 is shown with green dots. Scale: 10 µm. C: Quantification of PLA density expressed as number of dots per 100 µm2 (left), and PLA dots, expressed as number/cell (right). Data of 45 (WT), 20 (INF2-KO), 50 (INF2-KO + INF2-caax) and 61 (INF2KO + INF2-noncaax) from 2 independent experiments. Graph shows Mean ± S.D. One-way ANOVA used. **** p<0.0001. D: Western blot from GFP-trap pull-down assay, in HEK IP3R2 (R2) (R1/R3 DKO) INF2-KO cells transfected with GFP-INF2-caax or GFP-INF2-noncaax, in basal conditions (untreated, UT) or after 0.5 µM Latrunculin A treatment (Lat A). Elution (right) shows the proteins that Co-immunoprecipitated with INF2. E: Western blot images of candidate proteins from GFP-trap pull-down assay, in HEK IP3R2 expressing INF2-KO cells transfected with GFP-tagged INF2 constructs (WT and serine-mutants).

**Figure S5.**
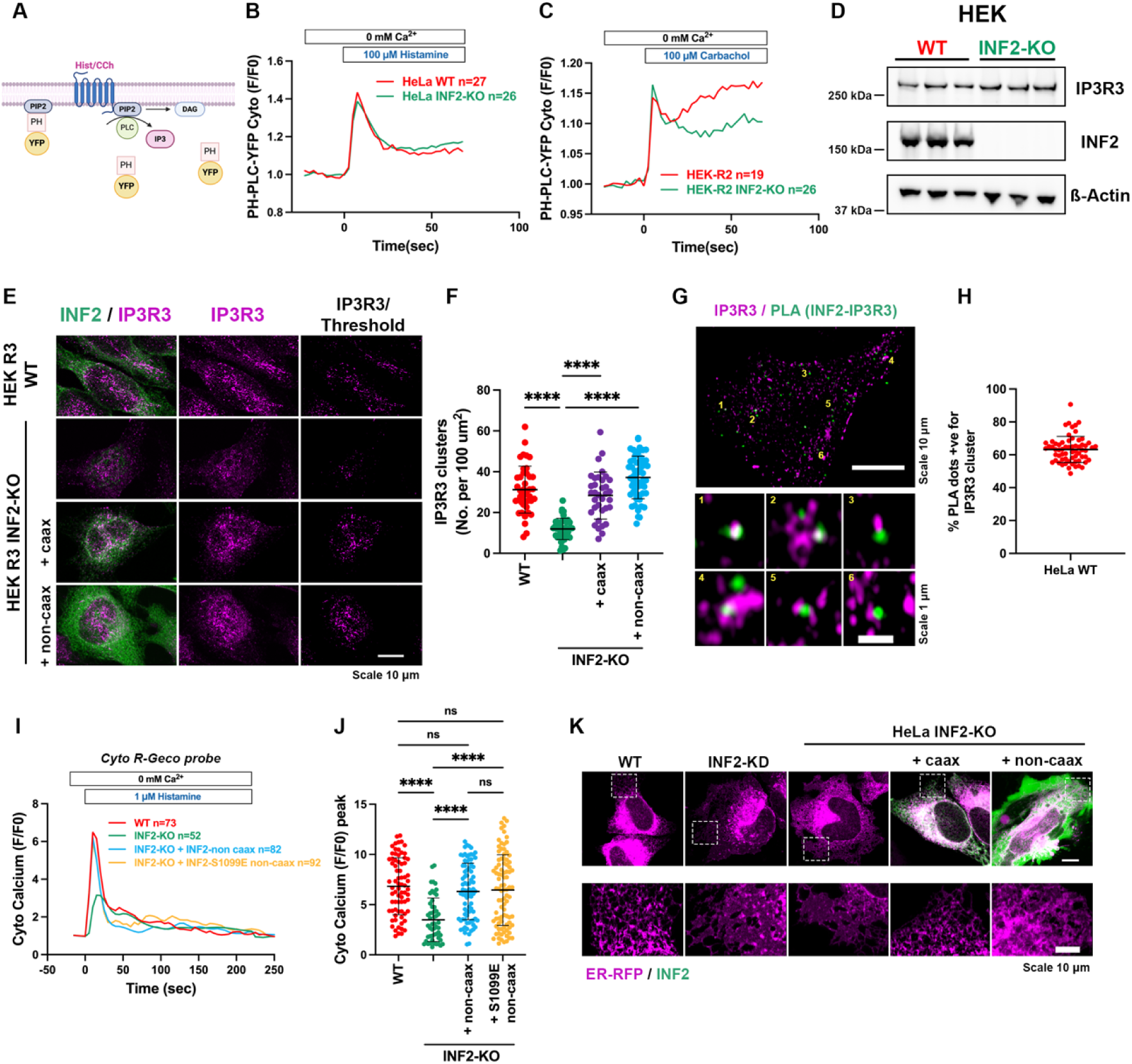
INF2 regulates formation of functionally competent IP3R clusters in cells. A: Diagram showing the YFP-PH-PLCδ construct, that bind to PIP2 in the plasma membrane. Agonist (histamine/carbachol) induced activation of GPCR pathway leads to phospholipase C mediated hydrolysis of PIP2 in the membrane forming mobile IP3 and membrane bound DAG. This caused re-localization of the probe in the cytosol. B: Fold change (F/F0) of the cytoplasmic YFP-PH-PLCδ fluorescence measured in WT and INF2-KO HeLa cells stimulated with 100 µM of Histamine. Data shown are average trace collected from 27 (WT) and 26 (INF2-KO) cells across 2 independent experiments. C: Fold change (F/F0) of the cytoplasmic YFP-PH-PLCδ fluorescence measured in WT and INF2-KO HEK-IP3R2 expressing (R1/R3 DKO) cells stimulated with 100 µM of carbachol. Data shown are average trace collected from 19 (WT) and 26 (INF2-KO) cells across 2 independent experiments. D: Western blot showing abundance of IP3R3 in HEK IP3R3 (R3) (R1/R2 DKO) WT or INF2-KO cells, β-actin used as loading control. E: Representative confocal micrographs of HEK-IP3R3 expressing cells (WT, INF2-KO and rescues), immuno-stained for INF2 (green) and IP3R3 (magenta). IP3R3 clusters after thresholding shown in the right-most panel. Scale: 10 µm. F: Quantification of IP3R3 clusters, expressed as number of clusters per 100 µm2 of cell area (from data set in E). Data from 42 (WT), 45 (INF2-KO), 35 (INF2-KO+INF2-caax) and 55 (INF2-KO+INF2-non-caax) cells from 2 independent experiments Graphs shows Mean ± S.D. One-way ANOVA used. **** p<0.0001. G: Confocal micrograph showing IP3R3 (magenta) and IP3R3-INF2 PLA dots (green) in HeLa WT cells. Inset shows partial co-localization between the PLA dots and IP3R3 clusters. Scale: 10 µm (main panel), 1 µm (inset) H: Quantification of the colocalization of PLA and IP3R3 dots, expressed as % PLA dots positive for IP3R3 cluster per cell. Data from 64 WT cells from 3 independent experiments. Graph shows Mean ± S.D. I: Cytosolic Ca2+ transients induced by 1 µM histamine in Cyto R-GECO transfected HeLa WT (red), INF2-KO (green), INF2-KO + INF2-noncaax WT (cyan), or INF2-noncaax-S1099E phospho-mimetic mutant (yellow). J: Quantification of maximal peak of Cyto Ca2+ from I, expressed as F/F0. Data from 73 (WT), 52 (INF2-KO), 82 (INF2-KO + INF2 noncaax), and 92 (INF2-KO + INF2-noncaax S1099E) cells across 2 independent experiments. Graphs show Mean ± S.D. One-way Anova used. **** p<0.0001 K: Confocal micrographs from WT and INF2 depleted (KD and KO) HeLa cells along with INF2-KO cells rescued with INF2 isoforms. The cells transfected with ER-RFP to delineate ER structure. Scale: 10 µm.

**Figure S6.**
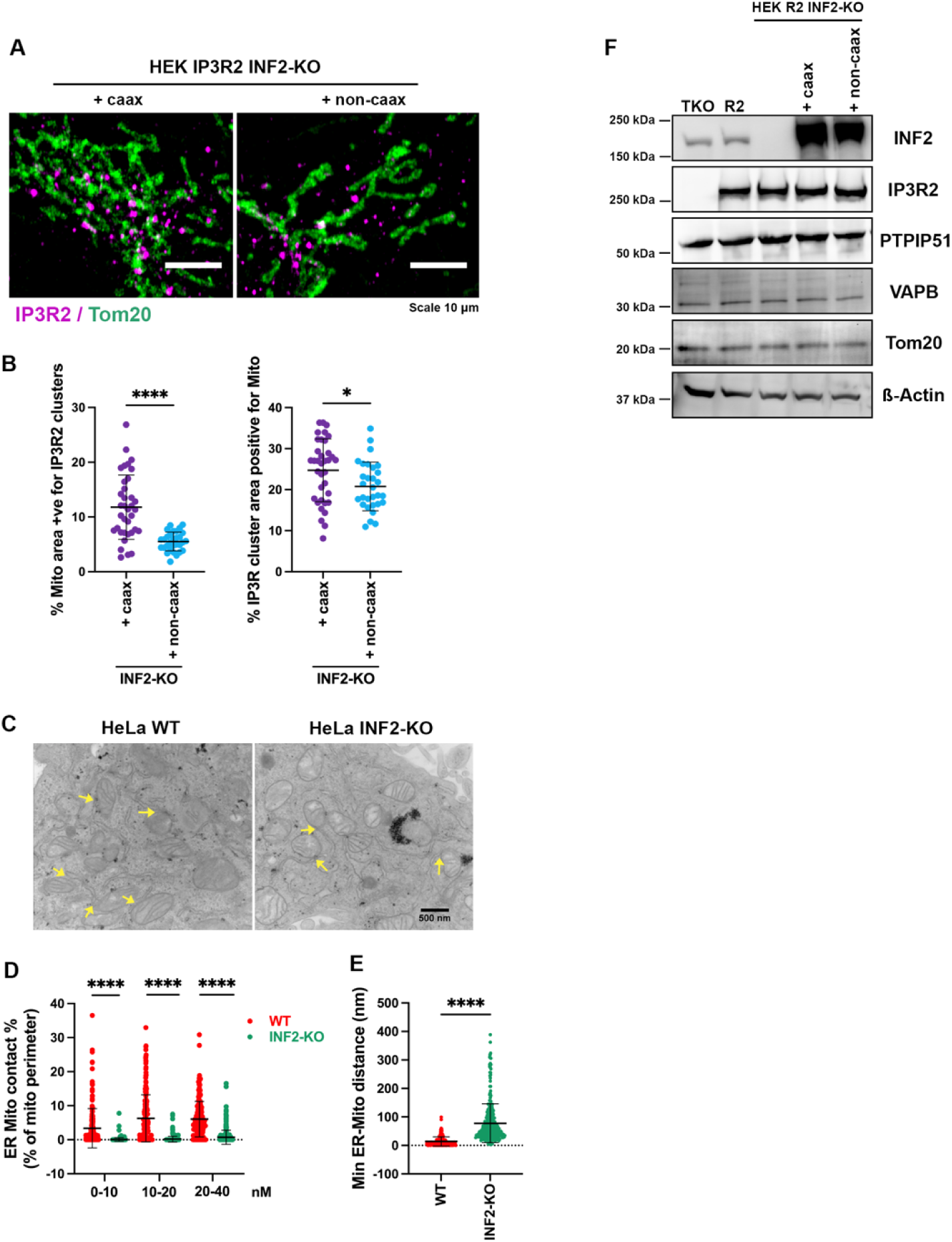
INF2-caax regulates ER-Mito contacts. A: Confocal micrographs of INF2 depleted IP3R2 expressing HEK-293 cells rescued with INF2-caax or INF2-noncaax. The cells were co-transfected with GFP-Tom20 to identify mitochondria. Cells were stained for endogenous IP3R2 (magenta). Scale: 10 µm. B: Quantification of IP3R2 clusters on the mitochondria expressed as % Mitochondria area positive for IP3R2 clusters or % IP3R2 cluster area positive for mitochondria per cell. Data from 36 (INF2-Caax) and 30 (INF2-noncaax) expressing cells across 2 independent experiments. Graphs shows Mean ± S.D. Student’s T-test used. **** p<0.0001. C. Representative images from thin-section TEM in HeLa WT and INF2-KO cells showing ER and mitochondria in the sections. The arrows represent close contact between ER and Mitochondria in the sections. Scale: 500 nm. D. Scatter plots showing the percentage of close ER-Mito contacts, expressed as % of mitochondrial perimeter, between 0-10, 10-20 and 20-40 nm. Data of 190 and 297 mitochondria quantified in WT and INF2-KO, respectively, from 2 independent sets of images. Graphs show Mean ± S.D. unpaired Student’s T-test. **** p<0.0001. E. Quantification of the minimal ER-Mito distance in nm. Data of 190 and 297 mitochondria quantified in WT and INF2-KO, respectively, from 2 independent sets of images. Graphs show Mean ± S.D. unpaired Student’s T-test. **** p<0.0001. F. Western blot showing the expression of INF2, IP3R2, VAP-B, PTPIP51 and Tom20 in HEK IP3R-TKO, IP3R2 (R1/R3 DKO) WT, INF2-KO and INF2-KO rescues cells. β-Actin was used as loading control.

